# *The molecular interplay of the establishment of an infection – gene expression of Diaphorina citri* gut and *Candidatus* Liberibacter asiaticus

**DOI:** 10.1101/2021.04.06.438581

**Authors:** Flavia de Moura Manoel Bento, Josiane Cecília Darolt, Bruna Laís Merlin, Leandro Penã, Nelson Arno Wulff, Fernando Luis Cônsoli

## Abstract

*Candidatus* Liberibacter asiaticus (CLas) is one the causative agents of greening disease in citrus, an unccurable, devastating disease of citrus worldwide. CLas is vectored by *Diaphorina citri*, and the understanding of the molecular interplay between vector and pathogen will provide additional basis for the development and implementation of successful management strategies. We focused in the molecular interplay occurring in the gut of the vector, a major barrier for CLas invasion and colonization. We investigated the differential expression of vector and CLas genes by analyzing a *de novo* reference metatranscriptome of the gut of adult psyllids fed of CLas-infected and healthy citrus plants for 1-2, 3-4 and 5-6 days. CLas regulates the immune response of the vector affecting the production of reactive species of oxygen and nitrogen, and the production of antimicrobial peptides. Moreover, CLas overexpressed *peroxiredoxin* in a protective manner. The major transcript involved in immune expression was related to melanization, a *CLIP-domain serine protease* we believe participates in the wounding of epithelial cells damaged during infection, which is supported by the down-regulation of *pangolin*. We also detected that CLas modulates the gut peristalsis of psyllids through the down-regulation of *titin*, reducing the elimination of CLas with faeces. The up-regulation of the neuromodulator *arylalkylamine N-acetyltransferase* implies CLas also interferes with the double brain-gut communication circuitry of the vector. CLas colonizes the gut by expressing two *Type IVb pilin flp* genes and several chaperones that can also function as adhesins. We hypothesized biofil formation occurs by the expression of the cold shock protein of CLas. We also describe the interplay during cell invasion and modification, and propose mechanisms CLas uses to invade the host hemocel. We identified several specific targets for the development of strategies directed to interfere with the successful utilization of the psyllid vector by this pathogen.

**Author Summary:** Huanglongbing (HLB) or greening is an incurable disease causing severe damage to citrus production, making citrus industrial activity unsustainable in several countries around the world. HLB is caused by three species of *Candidatus* Liberibacter. *Ca.* L. asiaticus (CLas), vectored by the psyllid *Diaphorina citri*, is the prevalent species. Attempts to apply new technologies in the development of strategies for disease and pest management are been made. However, we still miss basic information on this system to efficiently apply the current technologies and envisage the implementation of new approaches for pest control, despite the relevant scientific contribution available. One major gap is regarded to the molecular interplay between CLas and its vector. We focused our attention in the molecular interplay occurring at the first relevant interaction of CLas and *D. citri*, represented by the gut barrier. We report the transcriptional activity of CLas during the invasion and establishment of the infection in the gut of the vector, as well as the transcriptional activity of the vector in response to the infection. We identified several host genes that are targeted and regulated by CLas as well as several CLas genes that are promising targets for the application of new management strategies.

## 1. Introduction

Microbes often establish different interactions with metazoans depending on the evolutionary history shared with their hosts. There are several examples of benefitial associations with multicellular hosts resulting from distinct ecological interactions with their hosts and associated trophic levels. But a concurrent number of interactions are also known to be noxious to multicellular hosts due to the development of a range of pathologies (Baumann 2005, Webb et al. 2006, Masnfield et al. 2012, Quigley 2013, Lu et al. 2020)

Viruses are certainly the most common microbes infecting prokaryotes and eukaryotes (Fauci 2001, Weinbauer 2004), and the year long pandemics caused by the Sars-CoV2 virus is overshadowing other threatining pathogens to animals and plants. Bacteria are widely associated with multicellular organisms, causing devastating diseases. Bacteria-causing diseases in agricultural systems are a risk to food security, causing severe losses in animal and plant production (Mansfield et al. 2012, Abebe et al. 2020).

Bacterial plant pathogens are also adapted to infect and propagate in insect tissues, as many insects are used as vectors by plant pathogens to infect new host plants and spread themselves in the environment. Plant pathogenic bacteria vectored by insects that use persistent, circulative, propagative mode of transmission require plastic phenotypes to interact with the different enviroments represented by sieves and tissues of the host plant, and the diverse environments faced in the gut lumen, hemocele and different tissues of the vector insect during the processes of acquisition and vector competence development (Huang et al. 2020). The capability of these pathogens to infect completely different hosts (plant and insect), and yet to depend on the shuttle host (insect) to locate and collect the pathogen in a diseased plant, and later transport it to new, health host plants for the establishment of new infections requires a set of strategies devoted to manipulate the host insect (Perilla-Henao & Casteel 2016, Mauck et al. 2018). The understanding of the interactions pathogens and hosts have at their molecular level during the processes of host invasion and infection establishment is required for a fully comprehension of the mechanisms involved in guaranteeing the host – pathogen association. Such mechanisms of close interactions in the associations of host and pathogens represent potential new targets for the development of new technologies and/or use of existing technologies to interfere with the successful association pathogens establish with their hosts.

*Candidatus* Liberibacter are bacteria posing serious threat to food security worldwide by causing diseases to potatoes and citrus. Citrus is certainly the most severely damaged crop by *Ca.* Liberibacter infections, as three different species are known to infect citrus plants (*Ca.* L. asiaticus, *Ca.* L. americanus and *Ca.* L. africanus), causing the incurable greening or huanglongbing (HLB) disease (Bové, 2006; Graça et al., 2016; Tomaseto et al., 2019; Bassanezi et al., 2020). Candidatus Liberibacter did not make to the top 10 plant pathogenic bacteria (Masnfield et al. 2012), but HLB has caused a tremendous impact in the citrus industry and led to the complete elimination of citrus orchards and the interruption of citrus production in what used to be highly productive citrus centers (Dala-Paula et al. 2019, Bassanezi et al. 2020, Singerman & Rogers 2020).

*Candidatus* Liberibacter asiaticus (CLas) is the most spread in the world, currently infecting citrus in the major producing areas (CABI/EPPO 2017, Ajene et al. 2020, Wulff et al. 2020). In plants, *Candidatus* Liberibacter reside exclusively in the phloem sieve tubes (Jagoueix et al., 1994; Bové & Garnier, 2003; Bové, 2006; Bendix & Lewis, 2018), but it will infect several tissues of their host vector insect (Ammar et al., 2011a; Ammar et al., 2017; Kruse et al., 2017; Ammar et al., 2019). CLas is vectored by the worldwidely distributed Asian Citrus Psyllid (ACP) *Diaphorina citri* Kuwayama (Hemiptera: Liviidae) (Bové, 2006).

Despite the importance of this disease to the worldwide citriculture and all of the investments and efforts of the scientific community to understand the interactions of the pathogen with its vector, much of the mechanisms involved in the pathogen-vector interplay remain unknown. We learned that CLas is transmitted from host plant to host plant by establishing a persistent, propagative transmission mode with its vector, a transmission mode that is very common to plant viruses vectored by aphids and whiteflies (Gray et al., 2014; Ammar et al., 2016; Kruse et al., 2017). We also learned CLas infects salivary glands and the midgut of vectors, tissues that are often recognized as natural barriers to circulative, propagative pathogens as they can prevent translocation of pathogens within the vector host (Ammar et al., 2011 a; Ammar et al., 2011 b; Ammar et al., 2016; Ammar et al., 2020). More recently, new information on the higher efficiency of adults than psyllid nymphs as vectors of CLas has been reported, with adults displaying a higher successful rate of infection of healthy citrus plants than nymphs (Ammar et al. 2020). Nevertheless, little information at the physiological and molecular level on the interface of the interactions of CLas with key vector tissues is available (Molki et al., 2019).

The limitations in the availability of efficient, cost-effective strategies for the management of the vector and the disease prompted a large number of investigations for the exploitation of new technologies, and promising results were mainly obtained with RNAi-based approaches (Santos-Ortega & Killiny, 2018; Lu et al., 2019a; Lu et al., 2019b; Yu & Killiny, 2020).

In here we focused on investigating the metatranscriptome of the gut of adults of *D. citri* feeding on CLas-infected citrus plants for different periods of time in order to identify genes of CLas that are expressed during the colonization of the gut of adults and the differential gene expression in the gut of psyllids over time against psyllids feeding on healthy, CLas-free citrus plants. Our major goal was to understand the dynamics of CLas gene expression during psyllid colonization and the response mechanisms that were activated in the gut epithelium of psyllids when exposed to *Ca*. Liberibacter asiaticus cells. We believe our data represents a source of very specific targets for the development and implementation of new strategies of psyllid/disease control using RNAi and/or gene editing technologies such as the CRISPR-Cas9 system.

## 2. Results

### 2.1. *De novo* transcriptome assembly

Sequencing from libraries of the gut of uninfected and CLas-infected nymphs and adults of *D. citri* yielded 395,151,161 reads, with an average of 16,464,632 reads/library. The use of the resulting 385,677,949 trimmed, quality-filtered reads (average 16,069,915 reads/library) allowed the *de novo* assembly of a transcriptome with 260,612,776 nucleotides (260 Mb), with transcripts with an average size of 481 bp and a N50 of 2,095 bp. The assembly resulted in 248,850 transcripts with an average of 39.9% GC content. Annotation of the transcriptome allowed the putative identification of 90,531 transcripts (36.4%), of which 52,081 transcripts were allocated to different gene ontology categories (S1 Fig). Additional filtering using the highest score hit after BlastX allowed the identification of 66,993 transcripts belonging to *D. citri*, 807 belonging to *Ca.* Liberibacter, and 1967 to the secondary symbiont *Wolbachia*, among others (S2 Fig).

### 2.2. CLas gene expression in the gut of adults of *Diaphorina citri*

Since we could not detect relevant read countings against transcripts belonging to *Ca*. L. asiaticus when using samples obtained from nymphs, only samples collected at the adult stage were were subjected to differential expression analysis. Gene expression analysis recognized 807 transcripts belonging to *Ca*. L. asiaticus in the gut of adult psyllids fed on CLas-infected citrus plants in at least one of feeding exposure periods analyzed for adults (1-2 d; 3-4 d; 5-6 d), with no CLas-related transcripts being identified in gut samples from insects fed on healthy plants (*CLas*^−^).

The detection of gene expression of *Ca.* L. asiaticus in adult psyllids increased in adults feeding on CLas-infected plants for 1-2 d to 5-6 d. Thus, 725 transcripts of CLas were expressed in *A1CLas*^+^, 766 in *A2CLas*^+^ and 804 in *A3CLas*^+^ (Fig 1). One transcript was expressed only in *A1CLas*^+^ and *A2CLas*^+^, 22 in *A1CLas*^+^ and *A3CLas*^+^, and 63 in *A2CLas*^+^ and *A3CLas*^+^. We also identified CLas genes that were exclusively expressed at the early (1-2 d, *A1CLas*^+^= 1 transcript, DN14421_c0_g2_i2 = nicotinate-nucleotide adenylyltransferase), intermediate (3-4 d, *A2CLas*^+^= 1 transcript, DN18703_c1_g1_i24 = Flp type IVb pilin) and late stage (5-6 d, *A3CLas*^+^= 18 transcripts) after adult started feeding on CLas-infected citrus plants (Fig 1). Most of these transcripts were represented by several isoforms of a gene, with different isoforms expressed in more than one of the sampled feeding times. Only *A3CLas*^+^ adults had transcripts represented by unique isoforms of a gene specifically expressed in their gut (Table 1).

**Fig 1.**
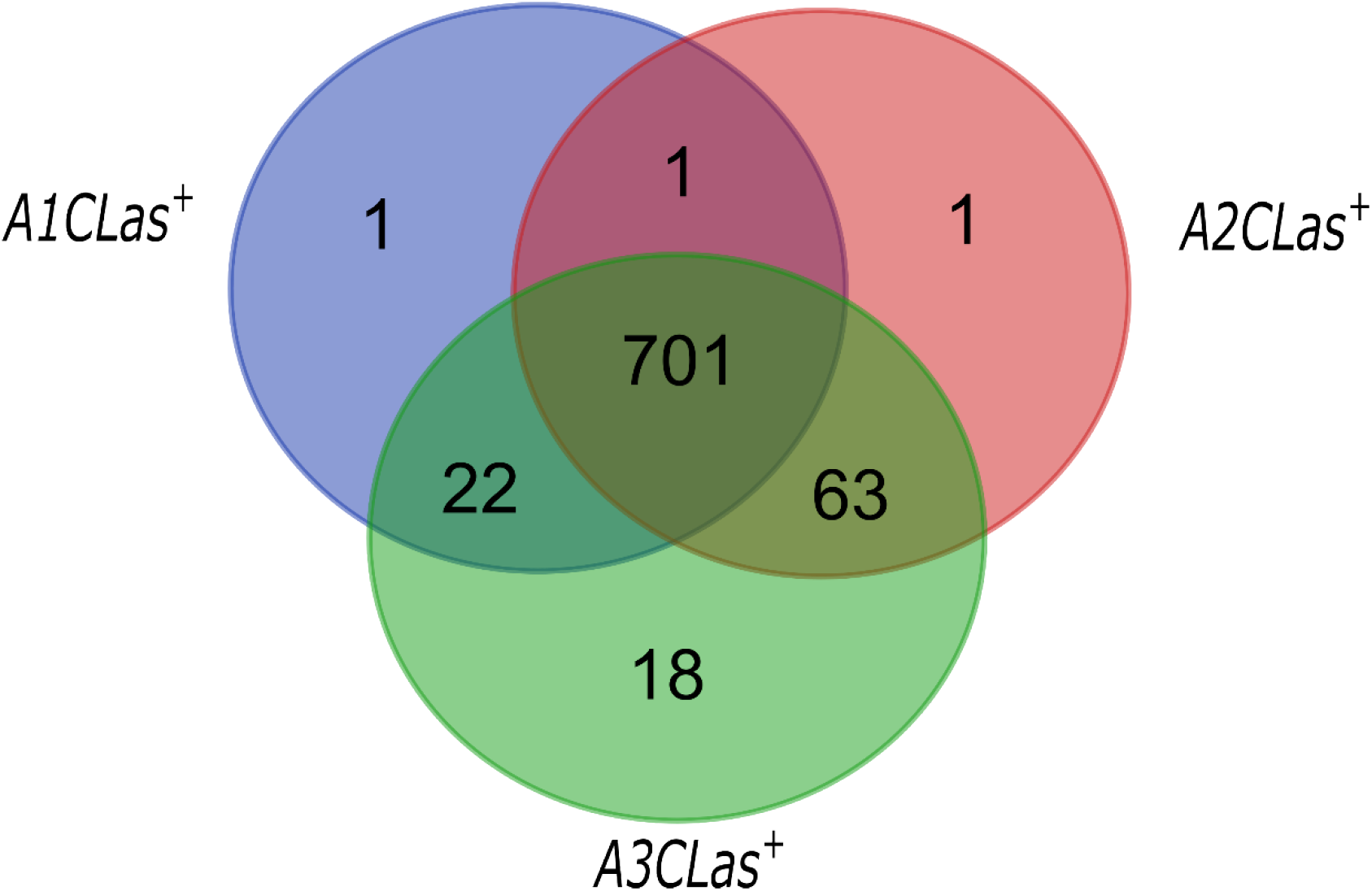
Venn diagram of *Candidatus* Liberibacter asiaticus transcripts differentially expressed in gut of adults at different periods of feeding on CLas-infected citrus plant. *A1CLas*^+^: adults that fed on CLas-infected citrus plant for 1-2 days; *A2CLas*^+^: adults that fed on CLas-infected citrus plant for 3-4 days; and *A3CLas*^+^: adults that fed on CLas-infected citrus plant for 5-6 days.

**Table 1.** Single isoforms of *Candidatus* Liberibacter asiaticus transcripts exclusively expressed in the gut of adult psyllids after 5-6d of feeding (*A3CLas*+) on CLas-infected citrus plants.

| Transcript ID | Annotation | TPM count |
| --- | --- | --- |
| DN1443_c0_g1_i1 | hypothetical protein PSGCA5_05 | 58.77 |
| DN15414_c0_g1_i2 | pyridoxine 5'-phosphate synthase | 5.33 |
| DN25987_c0_g1_i1 | hypothetical protein | 30.77 |
| DN32802_c0_g1_i1 | anti-repressor protein | 38.31 |
| DN39169_c0_g1_i1 | hypothetical protein BWK56_05555 | 69.16 |
| DN44737_c0_g1_i1 | threonine--tRNA ligase | 168.28 |
| DN47430_c0_g1_i1 | flagellar basal body rod protein FlgF | 54.15 |
| DN55271_c0_g1_i1 | phage related protein | 10.66 |
| DN62541_c0_g1_i1 | flagellar basal body P-ring protein FlgI | 31.40 |
| DN64078_c0_g1_i1 | putative phage-related acetyltransferase | 43.56 |

Pairwise gene expression analysis of CLas in the gut of adult psyllids after different periods of feeding on CLas-infected citrus plants revealed a high number of differentially expressed CLas genes in *A3CLas*^+^ as compared to *A1CLas*^+^ and *A2CLas*^+^ adults (S1 Table). One-hundred transcripts out of the over 700 transcripts detected in the three sampling times differed in their abundance (S1 Table). Differences in the level of expression were detected for 80 transcripts when comparing *A1CLas*^+^ and *A3CLas*^+^ and 20 transcripts in comparisons of *A2CLas*^+^ and *A3CLas*^+^ (S1 Table). CLas expression was always higher in *A3CLas*^+^ when compared to the others. No differences in gene expression between *A1CLas*^+^ and *A2CLas*^+^ were detected (S1 Table).

### 2.3. Differential gene expression in the gut of *CLas*^+^ and *CLas*^−^ adults of *Diaphorina citri*

Differential gene expression (DE) of the gut of adults of *D. citri* after feeding on CLas-infected plants for 1-2 days identified 24 DE transcripts in the gut of *CLas*^+^ insects, 20 of them were up-regulated and 4 were down-regulated in response to CLas infection. Most of the up-regulated transcripts (11) belong to uncharacterized proteins, while nine up-regulated transcripts were identified (Table 2). After 3-4 days of feeding on CLas-infected plants, the number of DE transcripts in the gut of *CLas*^+^ as compared to *CLas*^−^ adult psyllids increased to 35, most of which were up-regulated (27). Eighteen out of the 27 up-regulated transcripts, and seven out of the eight down-regulated transcripts were putatively identified (Table 2). The number of DE transcripts in the gut of adult psyllids after 5-6 days of feeding on CLas-infected plants was the highest, with 61 DE transcripts in the gut of *CLas*^+^ insects, 57 (33 unknown proteins) of them were up-regulated and 4 were down-regulated when compared to *CLas*^−^ adult psyllids (Table 2).

**Table 2.**
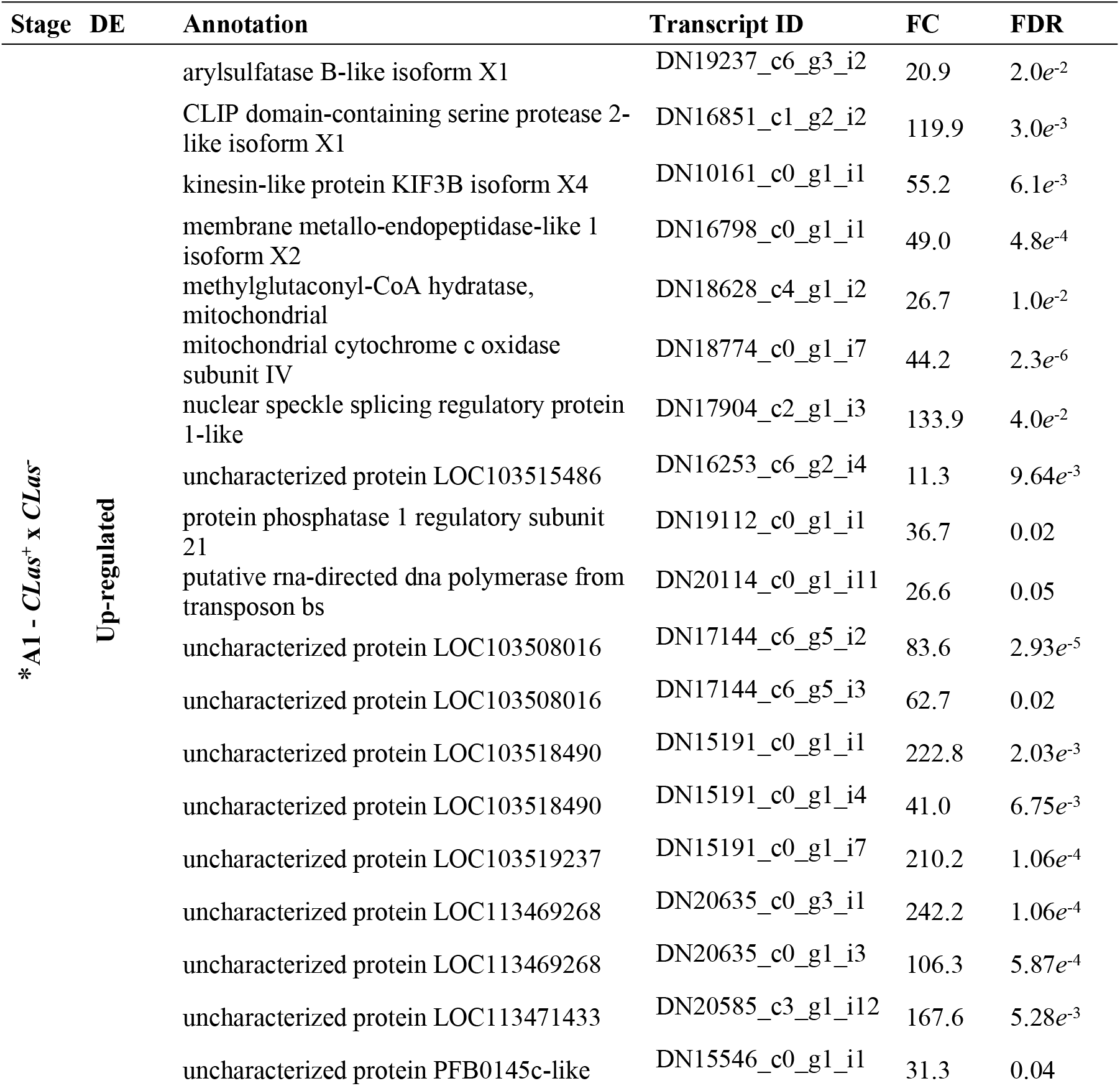
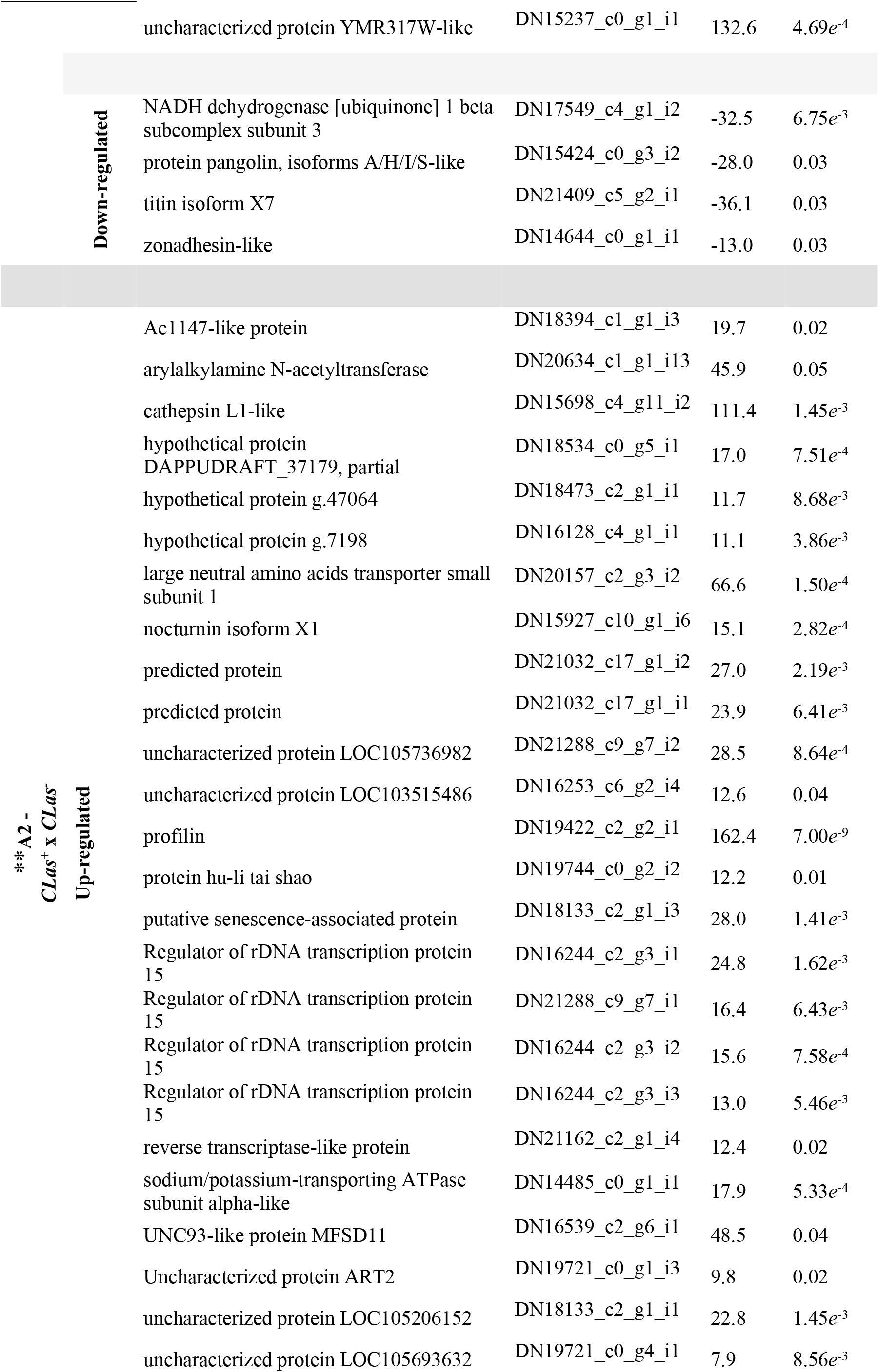
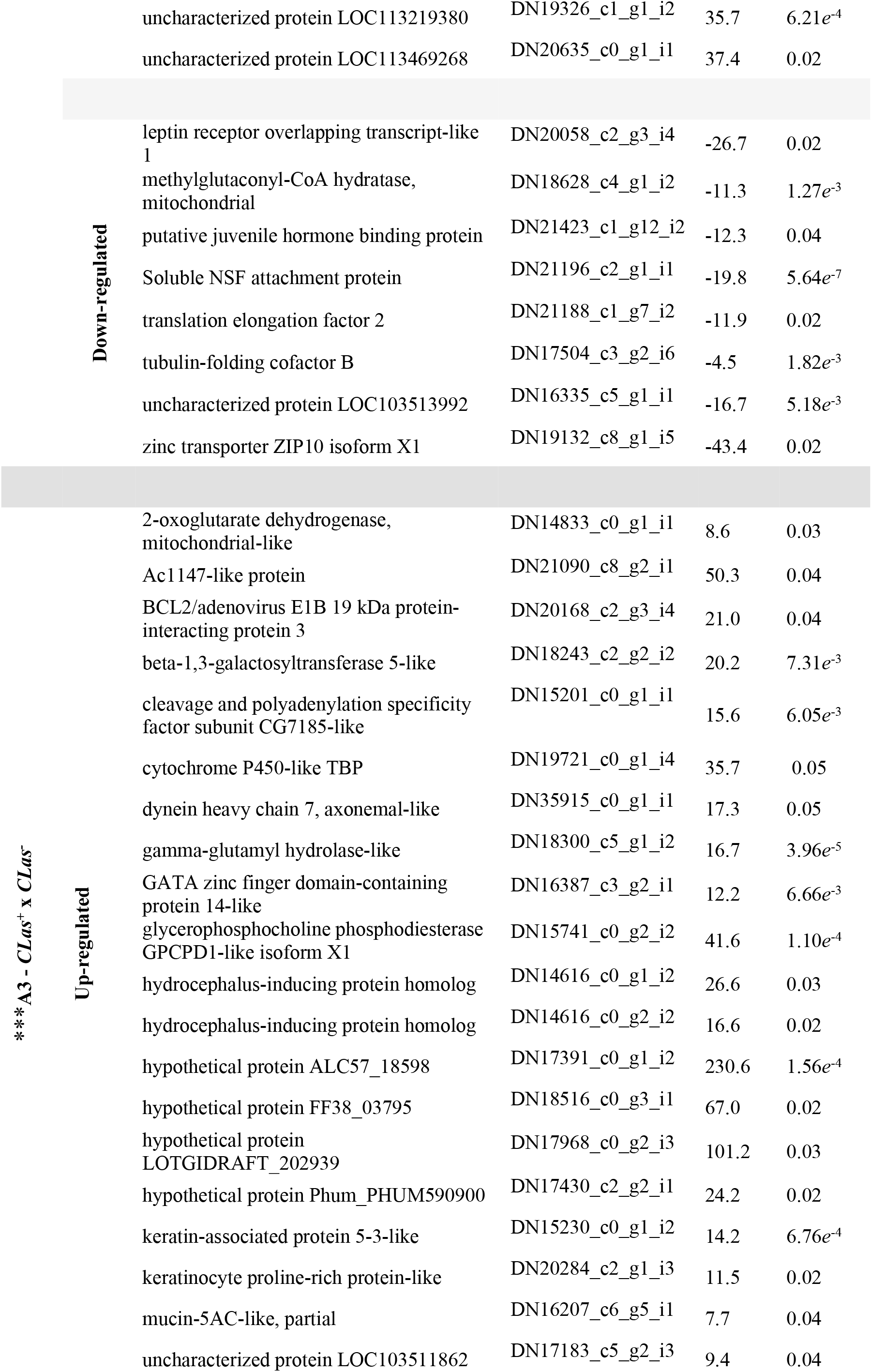
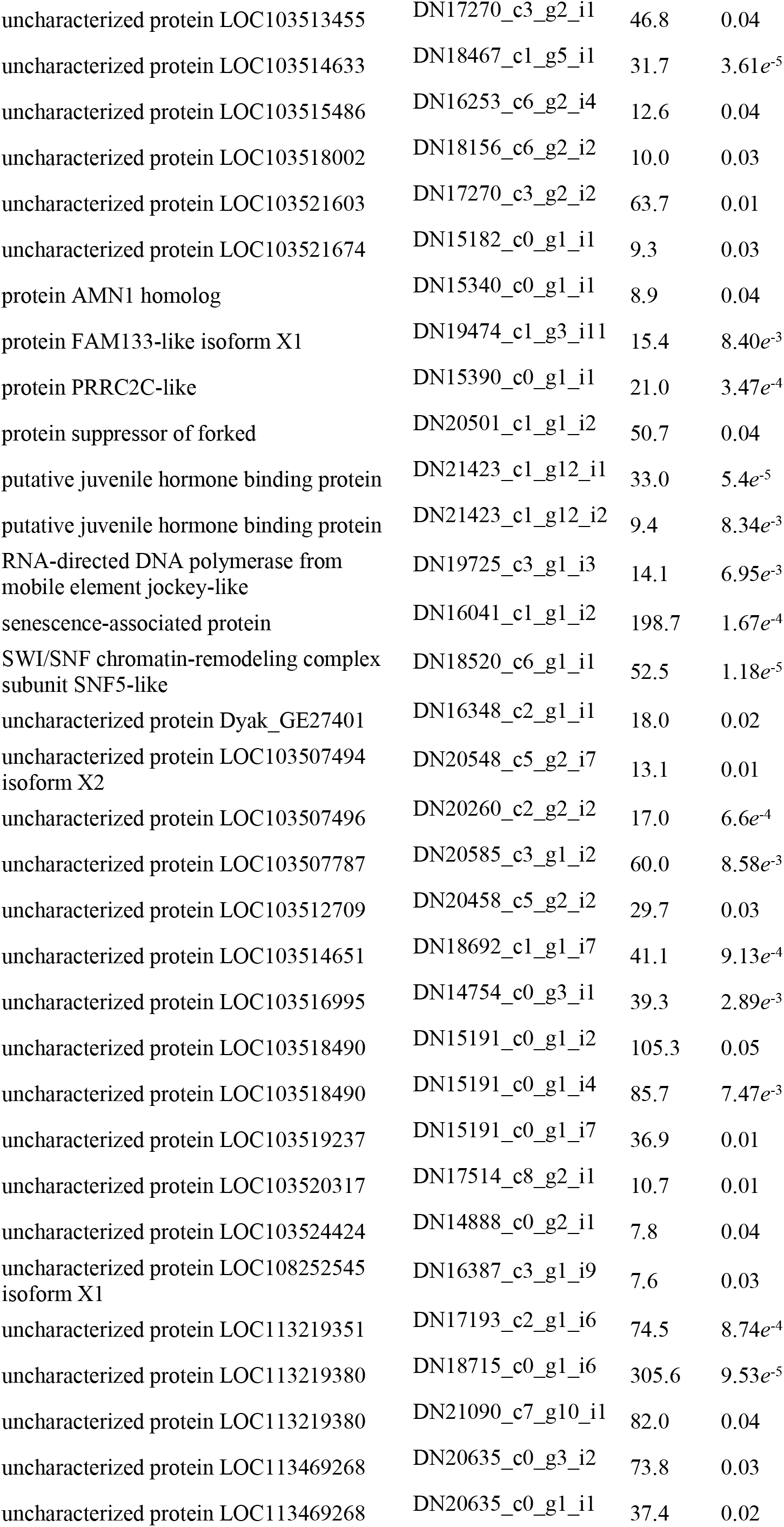
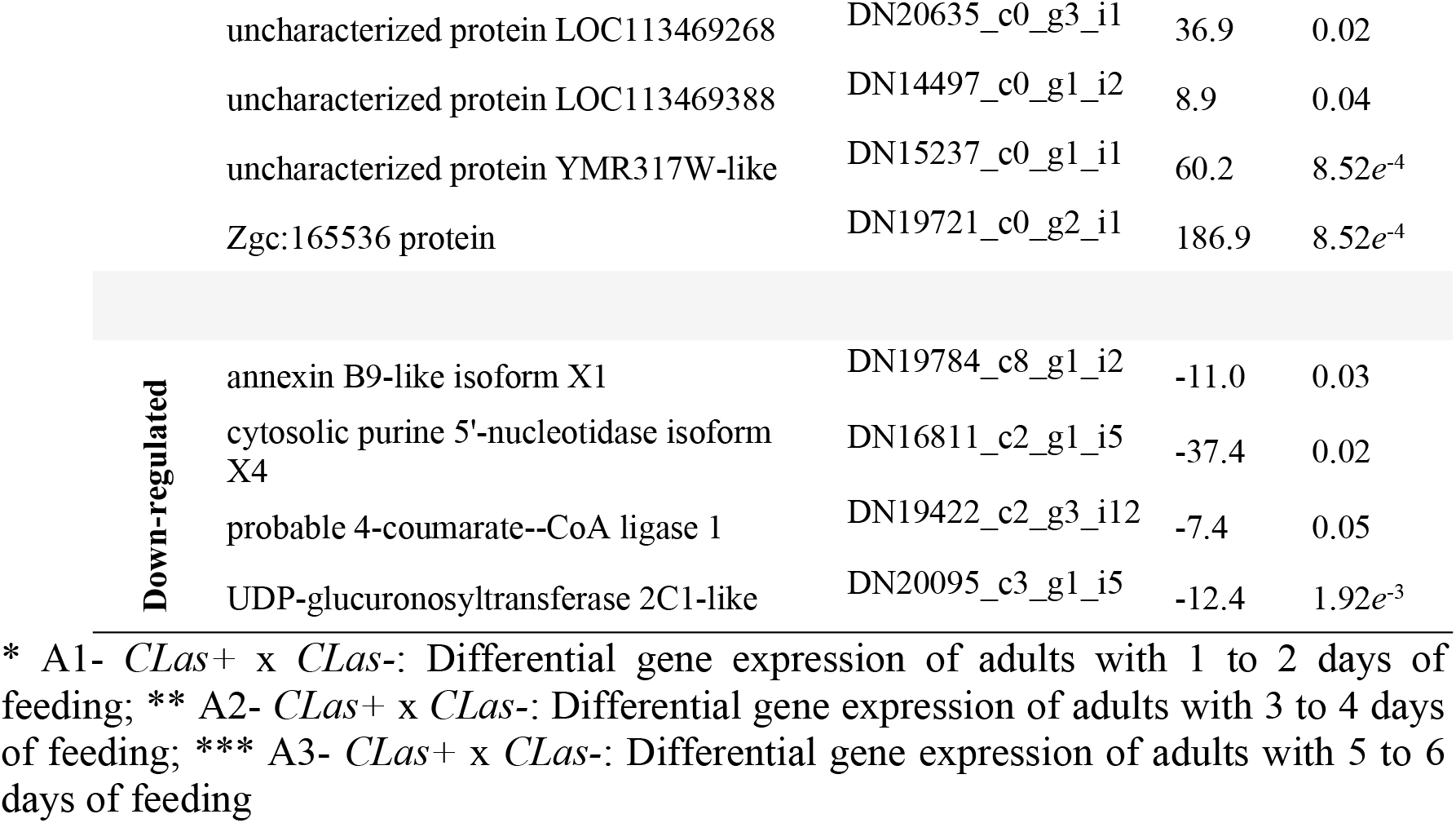
List of genes difference expressed comparison of *Diaphorina citri* adults fed on *Candidatus* Liberibacter asiaticus infected and non-infected plants at different

From the total number of differentially expressed transcripts of *CLas*^+^ when compared to their respective *CLas*^−^ psyllids, only one transcript was differentially expressed in *CLas*^+^ psyllids regardless the time they were allowed to feed on CLas-infected citrus plants as compared to psyllids fed on CLas-uninfected citrus plants (Fig 2). Most of the DE transcripts detected in *CLas*^+^ psyllids differed only at the specific stage they were compared. Seventeen (71%) of the DE transcripts detected in the gut of adult psyllids after 1-2 d, 31 (88%) after 3-4 d and 60 (88%) after 5-6 d of feeding in CLas-infected plants differed from controls specifically at each particular stage (Fig 2).

**Fig 2.**
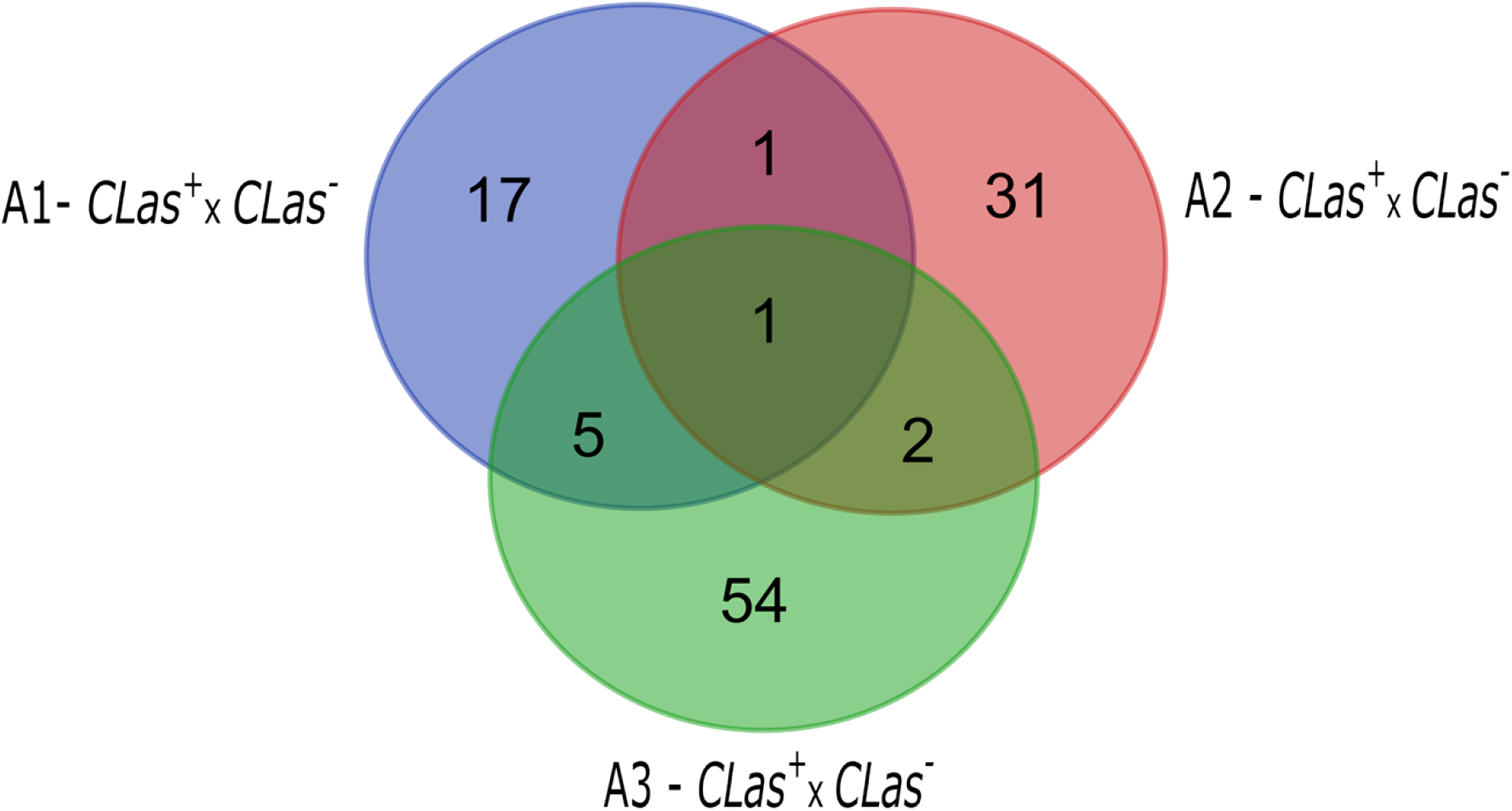
Venn diagram of *Diaphorina citri* transcripts differentially expressed in gut of adults at different periods of feeding on CLas-infected citrus plant compared to insects that fed on healthy citrus plant. A1 - *CLas*^+^ x *CLas*^−^: adults that fed on CLas-infected versus uninfected citrus plant for 1-2 days; A2 - *CLas^+^*x *CLas*^−^: adults that fed on CLas-infected versus uninfected for 3-4 days; and A3 - *CLas^+^*x *CLas*^−^: adults that fed on CLas-infected versus uninfected for 5-6 days.

## 3. Discussion

The gut differential gene expression in adult psyllids differed depending on the duration of feeding on CLas-infected citrus plants. Longer was the duration of feeding on CLas-infected citrus plants, higher were the number of DE transcripts. Most interesting, the majority of DE transcripts detected were specific to each one of the periods of feeding psyllids remained exposed to CLas-infected plants.

Gene expression of CLas in the gut of adult psyllids after different times of feeding on CLas-infected citrus plants were quite different. CLas transcription in the gut of adult psyllids was highly active soon after adult feeding started, but expression of a large set of genes was significantly increased at later stages of feeding (*A3CLas*^+^).

### Immune attack and immune defense responses

Bacteria that gain access to the hemocoel of insects find an easy way through the gut epithelium of the midgut, once the epithelium in this region does not have the cuticle lining protecting the fore- and the hindgut. Psyllids, as other sap-feeding insects do not carry the peritrophic membrane that protects the midgut epithelium of several other groups of insects (Lehane & Billingsley 1996, Erlandson et al. 2019). But regardless the presence of such physical barriers, the front line of defense of the gut immune system involves the activation of the epithelial innate immune system for the production of reactive oxygen species (ROS). Bacteria that survive in the gut of insects are either resistant to or are able to metabolize ROS (Buchon et al. 2013, Vallet-Gely et al. 2008).

Analysis of the CLas gene expression in the gut of adult psyllids demonstrated that CLas expressed the cytoprotective antioxidant enzyme peroxiredoxin (DN17546_c4_g1_i1), capable of reducing ROS and reactive nitrogen species (RNS) produced in the process of gut infection (Perkins et al., 2015; Knoops et al., 2016). The significant increase in the expression of peroxiredoxin from *A1CLas*^+^ to *A3CLas*^+^ (FC = 13.48) indicates this enzyme provides an increased contribution in the infection process as CLas cells also established an intracellular interaction with the gut epithelium of psyllids. CLas expression of peroxiredoxin occurred despite the expected reduction of ROS in the gut epithelium of adult psyllids, as the major source for ROS production in the gut, NADH dehydrogenase [ubiquinone] 1 beta subcomplex subunit 3 (DN17549_c4_g1_i2), was down-regulated in *CLas*^+^ psyllids.

These results suggest that CLas can inhibit hydrogen peroxide production in the host insect, and that peroxiredoxin is certainly serving roles other than its antioxidative contribution. The adipokinetic hormone (ADK) is involved in the regulation of the oxidative stress response in insects (Kodrík et al. 2015). Unfortunately, the expression levels of ADK-related genes by CLas infection could not be verified as ADK is produced and stored in neurosecretory cells of the *corpora cardiaca*. Nevertheless, regulation of this neuropeptide hormone in the gut epithelium of *CLas*^+^ psyllids could occur by the up-regulation of the metallo-endopeptidase-like 1 isoform X2 (DN16798_c0_g1_i1) neprilysin. Neprilysins are metalloproteases better known for their neuropeptide degrading activity, while also carrying peptide-degrading activity in other tissues, including the gut (Turner et al. 2001).

CLas peroxiredoxin could also be acting in the regulation of H_2_O_2_-mediated cell signaling processes, such as those involved in growth and immune responses (Shears & Hayakawa 2019). The oxidative response produced by gut epithelia due to microbial infection leads to cell proliferation and modulation of the innate immune response (Neish 2013), and ROS is required for inducing nitric oxide production in the gut. Nitric oxide production will in turn trigger the synthesis of antimicrobial peptides (AMP) and activate organ-to-organ communication (Wu et al. 2012). Therefore, the down-regulation of ROS production in the gut of *CLas*^+^ psyllids could explain the lack of differential expression of genes belonging to the AMP production pathways. AMP synthesis can also be elicited by the recognition of peptidoglycans released by bacteria by membrane-bound peptidoglycan recognition proteins if peptidoglycans survive to amidase degradation (Vallet-Gely et al. 2008).

There is very little information on insect gut phagocytosis, although proteomic analysis of anal droplets of the beetle *Cryptorhynchus lapathi* larvae led to the identification of several proteins to support an immune cellular response at the gut level (Jing et al. 2018). Moreover, phagocytic receptors identified in the gut epithelium of *Drosophila* were proven to play crucial role in the phagocytosis of both Gram bacteria, and in controlling hemocele infection by bacteria invasion through the gut epithelium (Melcarne et al. 2019). We could not detect any changes at the molecular level in *CLas*^+^ psyllids that would demonstrate a phagocytic response in the gut, as well as we did not observe adult psyllids to build an immune response to CLas infection by activating ROS and AMP pathways.

Nevertheless, differential gene expression analysis of the gut of *A1CLas*^+^ psyllids demonstrated the mounting of a defensive response of the gut epithelium based on the increased expression of a CLIP domain-containing serine protease 2-like (DN16851_c1_g2_i2). CLIP domain-containing serine protease 2-like has been recently reported to be up-regulated in the midgut of CLas-infected psyllids (Yu et al., 2020). Serine proteases containing a CLIP domain are involved in the regulation of humoral responses through the activation of prophenoloxidases (PPO) and the Toll immune signaling pathway. Activation of Toll pathway leads to antimicrobial peptides production, while the activation of PPO results in melanogenesis (Anderson 2000; Nakhleh et al. 2017). Toll activation requires a multi-step proteolytic cascade (Stokes et al. 2015), and the lack of additional differentially expressed serine proteases and the antimicrobial peptides transcripts in the psyllid transcriptome support the argument that the up-regulation of CLIP-domain serine protease in the gut of ACP feeding on CLas-infected plants leads to the activation of melanization as an immune response against CLas infection. We also believe the activation of melanization at this stage of psyllid-CLas interaction was triggered by a local response caused by cell injury and invasion of the gut epithelium by CLas cells, resulting in the production of wound clots in order to avoid infected, damaged cells to suffer further damage, die and be replaced by *nidi* cells. This hypothesis is also supported by the down-regulation of pangolin (*Pan*) (DN15424_c0_g3_i2), a key regulator of the Wnt/Wg pathway. Wnt proteins are highly conserved and participate in the control of growth, patterning, tissue and energy homeostasis. Pangolin was reported to bind to the β-catenin homologue Armadillo, acting directly in the regulation of gene expression in response to Wnt signaling (Brunner et al. 1997; Franz et al. 2017). The Wnt signaling in *Drosophila* is an important process for the homeostasis of the gut tissue as it is involved in the regeneration of adult gut epithelial cells (Strand & Micchelli 2011, Cordero et al. 2012). Thus, the down-regulation of pangolin interferes with the activation of the Wnt signaling to regulate gene expression involved in the replacement of CLas-infected cells by the activation of the nidi cells to differentiate into new, active cells of the gut epithelium.

Studies on the role of genotype-genotype interactions of insects and parasites demonstrated activation and differential gene expression of the host immune machinery depending on the interacting parasite genotype. Alternative splicing was also reported as a required element in the specificity of the immune response of insects to specific parasite genotypes (Riddell et al. 2014). The nuclear speckles carry pre-mRNA splicing machinery composed by nuclear speckle-related proteins that mediate alternative splice site selection in targeted pre-mRNAs (Girard et al. 2012; Galganski et al., 2017). The detected up-regulation of proteins that mediate alternative splice site selection in targeted mRNAs, the nuclear speckle splicing regulatory protein 1-like (DN17904_c2_g1_i3) in *CLas*^+^ psyllids, suggests CLas acts on the regulation of gene expression of infected psyllids at the molecular level.

The psyllid initial defense response against CLas infection also included the up-regulation of arylsulfatase B (DN19237_c6_g3_i2), an enzyme stored in lysosomes that acts on large glycosaminoglycan molecules by removing attached sulfate groups (Bhattacharyya et al., 2014). Glycosaminoglycans are components of proteoglycans commonly exploited by pathogens as receptors for their adherence to different tissues (García et al. 2016). Glycosaminoglycans are produced by some pathogenic bacteria as extracellular capsule and used in the process of host infection and colonization to facilitate pathogen attachment, invasion and/or evasion of host defensive mechanisms (Roberts, 1996). Arylsulfatases B can also participate in processes of biotransformation, mediating the sulfonation of xenobiotics by other enzymes to facilitate their excretion (Zhao et al., 2016). In this case, increase in the arylsulfatase could be related to the psyllid physiological needs to metabolize the high levels of flavonoids (hesperidin, narirutin and dydimin) produced in citrus plants infected by *Ca.* Liberibacter (Massenti et al., 2016; Dala-Paula et al., 2018; Kiefl et al., 2018). Flavanoids are important metabolites produced by plants in response to abiotic and biotic stressors and play a relevant contribution in plant resistance against microbes and herbivores (Treutter 2006), including citrus plants in response to CLas infection (Hijaz et al. 2013, 2020).

The overexpression of immune related genes and the required increased cell metabolism early in the process of interaction of the psyllid gut epithelium with CLas can explain the up-regulation of the mitochondrial cytochrome c oxidase subunit IV (DN18774_c0_g1_i7) in *CLas*^+^ insects. Mitochondrial cytochrome c oxidase subunit IV is the major regulation site for oxidative phosphorylation by catalyzing the final step of electron transfer in mitochondria (Li et al. 2006b).

### Cross-talk in the colonization of the gut lumen

Bacteria that survive the harsh chemical environment and the immune barriers available in the gut will have to use strategies to avoid their rapid elimination from the gut with faeces (Vallet-Gely et al. 2008, Benguettat et al. 2018). Insects are active feeders and food fastly transits through the gut. The fast food transit in the gut has been argued as one condition to explain the controversial lack of a resident microbiome in lepidopteran larvae, for example (Hammer et al. 2017).

CLas seems to employ different strategies to colonize the gut lumen by avoiding its elimination with faeces: regulation of gut peristalsis, synthesis of adherence proteins, and biofilm formation.

CLas regulation of the psyllid gut peristalsis is thought to occur through the up-regulation of the host neprilysin as earlier discussed. Neprilysins are reported to degrade peptide hormones, and the gut of insects contains several hormone-producing cells. Peptide hormones produced in the gut act as signaling molecules that are involved in the regulation of a range of processes, including gut peristalsis (Wegener & Veenstra 2015, Caccia et al. 2019, Wu et al. 2020). The hypothesis that CLas modulates the gut peristalsis of psyllids is supported by the down-regulation of titin (DN21409_c5_g2_i1) transcription in *CLas*^+^ psyllids. Muscle degeneration is marked by a reduction in titin proteins, and gut-associated muscles are in charge of producing the peristaltic movement observed in the gut. Additionaly, the intracellular non-catalytic domain of neprilysins in excess is shown to induce muscle degeneration (Panz et al. 2012). Thus, the degeneration of muscles associated with the gut would certainly impair gut peristalsis and consequently reduce the elimination of bacteria with faeces (Vallet-Gely et al. 2008, Benguettat et al. 2018).

Regulation of signaling in the gut is also evidenced by the up-regulation of arylalkylamine N-acetyltransferase (DN20634_c1_g1_i13) transcription in *A2CLas*^+^ psyllids. Arylalkylamine N-acetyltransferases are involved in the N-acetylation of arylalkylamines, playing an important role in the synthesis of melatonin in vertebrates and invertebrates (Klein 2007; Hiragaki et al. 2015). But in insects, arylalkylamine N-acetyltransferases also inactivate arylalkymines that play important roles as neuromodulators, such as octopamine, dopamine and serotonin (Brodbeck et al. 1998; Amherd et al. 2000). These neuromodulators are also implicated in the double brain-gut communication circuitry (Solari et al. 2017; Zhang et al. 2018).

CLas adherence to the psyllid gut was achieved by the expression of three genes of a subtype of the type IVb pilins, the tight adherence pili (Tad = Flp). Flp pilins are common to several Gram^+^ and Gram^−^ bacteria. Type IV pili are highly diverse and involved in a number of protein-protein interactions, but flp pili are better known for their role in adherence to living and nonliving surfaces. Pili are also involved in cell motility, secretion of exoproteins, and in host cell manipulation under extreme conditions (Giltner et al. 2012, Kazmierczak et al. 2015).

The expression of Type IVc pili of *Ca.* L. asiaticus was reported to be higher in psyllids than in plants, but only one (*flp3 pilin*) out of five *flp* genes was reported to be up-regulated in psyllids (Andrade & Wang 2019). The *flp3 pilin* gene was demonstrated to be under the regulation of VisN and VisR, demonstrating these proteins are important in the colonization of the host vector (Andrade & Wang 2019). In our analysis of the transcriptional profile of CLas in the gut of adult psyllids, we detected the expression of several isoforms of two *flp* genes (DN14425_c0_g1; DN18703_c0_g1), but both belonging to the family Type IVb pilin. Three isoforms of gene DN14425_c0_g1 (DN14425_c0_g1_i2; DN14425_c0_g1_i3; DN14425_c0_g1_i4) and two of gene DN18703_c0_g1 (DN18703_c0_g1_i21 and DN18703_c0_g1_i23) were highly and consistently expressed in the gut of psyllids in all sampling periods. Three LuxR family transcriptional regulators (DN1177_c0_g1_i1, DN14940_c0_g2_i1, and DN38633_c0_g1_i1) were also observed in the gut of *CLas*^+^ psyllids; nevertheless, none of them was identified as the VisR or VisN regulators of *flp3 pilin* gene reported by Andrade & Wang (2019).

The expression of CLas flagellins (*FlgA*) (DN3624_c0_g2_i1 and DN40834_c0_g1_i1) and several genes involved in the assembling and functioning of the flagellar machinery (*FliE, FliF*, *FliG*, *FliK, FliN, FliR, FliP, FlgB, FlgD, FlgE, FlgF, FlgG, FlgH, FlgI, FlgK, FlhA, FlhB, MotB,* and the *flagellar C-ring protein*) would indicate CLas assembles flagella for colonizing the gut of adult psyllids, although FlgF and FlgI were detected exclusively in *A3CLas*^+^. The expression of genes involved in the flagellar machinery corroborates recent data on the higher expression of genes encoding the flagellum apparatus of CLas in the gut of psyllids than in plants (Andrade et al. 2020a). Although we detected a much higher number of transcripts of the flagellar system of the CLas isolate we worked with than those reported to be expressed in the gut of psyllids by Andrade et al. (2020a), we did not detect the expression of three genes they evaluated (*fliQ, fliL*, and *flgA*). Differences in the overall number of genes of the flagellar system among lineages are expected to occur, but not among isolates of the same species (Bardy et al. 2003).

The production of flagellum in the gut by infecting CLas is also supported by transmission electron microscopy images, although most of the CLas cells observed lacked a flagellum (Andrade et al. 2020a). Analysis of images of the flagellum of CLas cells clearly shows CLas carries the secondary flagellar system, producing a lateral flagellum. The secondary flagellar systems arose twice in the evolution of bacteria, once in alpha-proteobacteria and once in the common ancestor of beta/gamma-proteobacteria (Liu & Ochman 2007). The primary flagellar system produces a polar flagellum and contributes with cell motility in liquid media, while the secondary flagellum system produces a lateral flagellum that is involved in adhesion and cell swarming on surfaces. Bacterial flagella serve bacteria not only as a motor apparatus, but also as a protein export/assembly apparatus. The motor apparatus of flagella can provide bacteria the movement required to remain in the gut, as demonstrated in the trypanosomatid *Vickermania* (Kostygov et al. 2020). Moreover, the flagellum pattern can be altered by bacteria depending on the environmental conditions faced, and bacterial flagella are recognized as important virulence factors (Moens & Vanderleyden 1996, Macnab 2003, Duan et al. 2013). Chemotaxis is an important contribution of bacterial flagellum to virulence (Matilla & Krell 2018), and the detection of the expression of four chemotaxis protein genes (DN15199, DN31800, DN46197 and DN50630) suggests the flagellum participates in cell adhesion to and swarming on the psyllid gut epithelium. Yet, we also propose the flagellum aids CLas to become chemically oriented for the localization of suitable host cells for invasion.

Biofilm formation requires bacteria to synthesize an extracellular matrix. Biofilms include extracellular proteins, cell surface adhesins and subunits of flagella and pili proteins (Fong & Yildiz 2015). We did not identify the expression of adhesins in the transcriptome of CLas in the gut of adult psyllids. Nevertheless, several chaperones were highly expressed, and many chaperones can act as adhesins in bacteria. Additionally, we believe the cold shock protein (DN10086_c0_g2_i1) of CLas is also playing key roles in biofilm formation. Cold shock proteins were demonstrated not only to participate in biofilm formation but also to support cell adhesion and motility, and to stimulate cell aggregation, interfering thus with virulence (Eshwar et al. 2017, Ray et al. 2020).

### Cross-talk for infecting the gut epithelium

Several gut-associated bacterial symbionts of hemipterans enter in close contact with epithelial cells to establish an intracellular phase through endocytosis (Nardi et al. 2019). CLas also enters the gut epithelium of psyllids through endocytosis, remaining inside vacuoles formed from or surrounded by endoplasmic reticulum membrane (Ghanim et al. 2017). The differential gene expression analysis of the gut of *CLas*^+^ psyllids led to the identification of transcripts involved with ultrastructural alterations of the gut epithelial cells, allowing the identification of the molecular intermediates that participate in the route CLas follows to invade the gut epithelium.

In addition to the regulation of the immune related responses earlier described in the gut epithelium of *CLas*^+^ psyllids, epithelial cells of the gut were altered early in the infection process (*A1CLas*^+^ adults). Alterations are perceived by the transcriptional down-regulation of zonadhesin-like (DN14644_c0_g1_i1), a transcriptional response already reported by Ramsey et al. (2015). Zonadhesin-like proteins represent an expansion of the zonadhesin multi-domain proteins that are implicated in the binding of sperm and egg in a species-specific manner. In the gut, zonadhesin-like proteins are thought to function as mucins, and two zonadhesin-like proteins of *D. citri* were previously predicted to be part of the extracellular matrix and to have a role in cell-cell adhesion. Mucins protect the gut epithelium from microbial infections and inflammation (Hansson 2012). Mucins are components of the perimicrovillar layer that in psyllids and other paraneopterans (Silva et al. 2004) replaces the chitin-based matrix (peritrophic membrane) that provides a protective barrier to the gut epithelial cells from pathogenic bacteria (Kuraishi et al. 2011).

In vertebrates the mucin composition of the mucous layer can vary among gut regions, with diet composition and gut microbial infection (Paone & Cani 2020). Mucins can also be exploited as a nutritional resource by bacteria, and degradation of mucins by a commensal microbe was demonstrated to facilitate the penetration of the epithelium by viruses (Schroeder 2019, Wu et al. 2019). The role different mucins can have on the host-pathogen interaction can explain the up-regulation of the psyllid gene coding for mucin-5AC (DN16207_c6_g5_i1) in *A3CLas*^+^ psyllids. The higher abundance of this protein in *CLas*^+^ psyllids has been demonstrated in previous proteomic analysis (Ramsey et al. 2017). Regulation of the host mucin-5AC has been reported in vertebrate hosts infected by Gram^−^ and Gram^*+*^ bacteria, and in one system modulation of mucin-5AC has been linked with increased adhesion of bacteria to the gut tissue (Dohrman et al. 1998, Quintana-Hayashi et al. 2015).

The differential expression of genes coding for cytoskeletal proteins involved in the regulation of the submembranous actin-spectrin network in cells of the gut epithelium of *CLas*^+^ psyllids demonstrates CLas interferes with the remodeling of gut epithelial junctions in adult psyllids, leading to endocytosis of intercellular junctions with cellular organelles. Adducins (protein Hu-li tai shao - DN19744_c0_g2_i2) form connections with membranes, promote spectrin-actin interactions and regulate actin filaments (Matsuoka et al. 2000). Internalization of apical junction and/or tight junction proteins may disrupt the epithelial barriers and favor the movement of bacterial toxins into cells (Hopkins et al. 2003).

The intracellular events in preparation for endocytosis can also be detected as indicated by the up-regulation of the psyllid phosphatase 1 regulatory subunit 21 (DN19112_c0_g1_i1) in the gut of *A1CLas*^+^. This protein participates in early (sorting process) or late (maturation) endosome pathway, which leads to the endocytosis of several types of materials, including pathogenic agents. Our hypothesis is that early endosomes will also internalize the leptin receptor overlapping protein-like 1. The leptin receptor overlapping transcript-like 1 (DN20058_c2_g3_i4) was down-regulated in the gut epithelium of *A2CLas*^+^ psyllids. Internalization of such receptors in early endosomes also affects cell signaling, and in this particular case can severely interferes with the gut epithelium resistance to microbial infection. Down-regulation of leptin/leptin receptor was reported to drastically affect cell resistance to amoeba and bacterial infection in several species (Faggioni et al. 2001, Guo et al. 2011, Mackey-Lawrence & Petri Jr 2012).

The endocytic vesicles detach and become free endocytic carrier vesicles transporting their cargoes to late endosomes to finally fuse with endoplasmic reticulum, where CLas cells aggregate in associated vacuoles as indicated by ultrastructural analysis (Ghanim et al. 2017). Fusion of lysosomes to cell aggregates vacuoles is inhibited by the down-regulation of soluble NSF attachment protein (SNAPs) (DN21196_c2_g1_i1) in *A2CLas*^+^. SNAPs are highly conserved proteins that participate in intracellular membrane fusion and vesicular trafficking (Stenbeck, 1998). Intracellular membrane fusion requires both SNAPs and NSF to act in concert, and inhibition of one of them will lead to the failure of membrane fusion and the accumulation of vesicles one cannot fuse (Rothman, 1994).

SNAPs are characterized by the presence of a tetratricopeptide repeat (TPR) domain (Lakhssassi et al., 2017). Tetratricopeptide repeat proteins (TRP) are directly involved in virulence, particularly due the translocation of virulence factors into host cells and in the blockage of phagolysosomal maturation, among others (Cerveny et al. 2013). Three TRPs (DN11192_c0_g1_i1; DN11192_c0_g1_i2; DN14826_c0_g1_i2) were expressed in the gut of *CLas*^+^ psyllids, but only DN14826_c0_g1_i2 was differentially expressed. The expression of different TRPs in all sampling periods and their time-specific differential expression indicates TRPs participates in different processes of CLas interactions with the host cells, from CLas establishment in the gut lumen to epithelial cell invasion and access to the hemocoel.

The down-regulation of the psyllids annexin B9-like isoform X1 (DN19784_c8_g1_i2) in the gut of *A3CLas*^+^ would interferes with the development of multivesicular bodies, once annexin B9 is involved in endosomal trafficking to multivesicular bodies (Tjota et al. 2011). This change will affect the transfer of the contents of CLas-containing endosomal vesicles to lysosomes for proteolytic degradation.

We propose that the establishment of the intracellular cycle and survival of CLas within host cells was putatively aided by the expression of CLas peptidyl-prolyl isomerase (PPI) (DN14386_c0_g1_i1), a protein that has been proved to participate in intracellular infection and virulence of other Gram^−^ bacteria (Norville et al. 2011, Pandey et al. 2017, Rasch et al. 2018). The Mpi PPI from *Legionella pneumophila* was demonstrated to require an active enzymatic site to enhance a proper host cell invasion (Helbig et al. 2003), although the preservation of the active site was not a requirement for the PPI of *Burkholderia pseudomallei* (Norville et al. 2011). Ghanim et al. (2017) suggested earlier that CLas cells do not enter the ER, but instead CLas cells recruit ER to transform the phagosome into a host-immune free space suitable for CLas survival and multiplication. These same ER-derived structures were already observed in *Legionella* and *Brucella* (Celli & Gorvel 2004, Robinson & Roy 2006).

Early endosomes and endoplasmic reticulum share contact sites that are used to bind to microtubules at or close to their contact sites, and both organelles remain bounded as endosomes traffic and mature (Friedman et al. 2013). Endosome trafficking within the cell involves the membrane binding to motor proteins and its transport along the actin and microtubule cytoskeleton (Granger et al. 2014). Trafficking of endosomes in *CLas*^+^ psyllids is provided by the up-regulation of kinesin-like protein KIF3B isoform X4 (DN10161_c0_g1_i1) in the gut of psyllids. Kinesins are motor proteins involved in the movement of multiple cytoplasmic organelles (Granger et al. 2014).

Profilin (DN19422_c2_g2_i1), another protein involved in intracellular movement of organelles, was also up-regulated in the gut of *A2CLas*^+^ psyllids. Profilins are actin-binding proteins capable of regulating actin polymerization and the availability of the actin cytoskeleton for binding to the endosomes; but new roles for profilins are being identified in vertebrates (Witke 2004). Additionally, profilin 1 play an important role in host–pathogen interactions. Intracellular pathogens such as *Listeria monocytogenes* and *Shigella flexneri* use the host-cell actin cytoskeleton to propel themselves through the cytoplasm and to spread to neighboring cells without entering the extracellular space (Kocks, 1994; Witke, 2004). Dynein (DN35915_c0_g1_i1), another motor protein, was up-regulated in the gut of *A3CLas*^+^ psyllids. Dyneins also contribute to microtubule-based transport in eukaryotic cells (reviewed in Holzbaur & Vallee, 1994; Porter, 1996; Hirokawa, 1998).

### Cross-talk for moving to the hemocoel

After CLas inhibits the early immune response and invades the intracellular space of the gut epithelium by regulating a clathrin-independent endocytosis mechanism, hiding from the host’s immune activators within vacuoles surrounded by endoplasmic reticulum membrane, where CLas multiplies, the molecular mechanism behind the release of CLas bacteria from epithelial cells in the hemocoel may require additional data.

The higher expression in *AD3CLas*^+^ of seven (DN15630_c0_g2_i3 = CLIBASIA_00255; DN15330_c0_g1_i2 = CLIBASIA_00880; DN14809_c0_g1_i = CLIBASIA_03170; DN14146_c0_g1_i1 = CLIBASIA = 04410; DN10141_c0_g1_i1 = CLIBASIA_00995; DN15141_c0_g1_i1 = CLIBASIA_01620; DN15414_c0_g1_i4 = CLIBASIA_01605) out of the 67 Sec-dependent proteins common to Las species (Thapa et al. 2020) indicates these candidate effector proteins play an important role in the dynamics of CLas infection of the psyllid gut epithelium at this stage. In fact, genomic analysis of Liberibacter species predicted a total of 166 proteins containing Sec-dependent signal peptides in the CLas strain psy62, from which 86 have been already experimentally validated (Prasad et al. 2016). These are potential effector proteins, and 106 of them share common homologues with Las_Ishi-1 and Las_gxpsy. But only 45 Sec-dependent proteins were shared among HLB-associated *Ca*. Liberibacter species (CLas, CLam and CLaf) (Wang et al. 2017; Andrade et al. 2020b). The detection of transcripts of the general secretion system provides further support for the participation of the detected Sec-dependent proteins in the late process of infection of the gut epithelium of *D. citri*. The Sec pathway secretion system transports proteins involved in bacterial cell functions and survival (Thapa et al. 2020). The Sec pathway is represented by two independent pathways, both detected in the gut of *CLas*^+^ psyllids: the post-translational pathway represented by the expression of SecA (DN14962_c0_g1_i1) and SecY (DN15568_c0_g3_i3); and the co-translational pathway represented by the signal recognition particle receptor FtsY (DN1686_c0_g1_i1) (Green & Mecsas 2016). We did not detect SecB expression in the gut of adult psyllids, indicating SecA can also act as a chaperone complementing the required activity of SecB as reported in *Escherichia coli* (McFarland et al. 1993).

Combined ultrastructure, proteomics and transcriptomics analysis of the gut epithelium of CLas-infected psyllids suggested CLas acquisition into the hemocoel would rely on the programmed cell death of CLas-infected epithelial gut cells, although CLas infection has little impact on the fitness of adult psyllids (Kruse et al. 2017). The low fitness impact of CLas infection to adult *D. citri* indicates CLas acquisition into the hemocoel would occur through an exocytosis process, as suggested for the *Ca.* Liberibacter solanacearum vectored by potato psyllids (Cicero et al. 2017).

By following the proposed subroutines based on mechanistic and essential aspects of cell death proposed by the Nomenclature Committee on Cell Death (Galluzzi et al 2018), our transcriptomic analysis could not support the proposition that CLas-infected epithelial gut cells of psyllids would initiate the process of programmed cell death (PCD) as proposed by Kruse et al. (2017). We did not observed transcription of caspases, a very common protein to several subroutines of cell death, as well as other aspects that would allow this characterization (Galluzzi et al 2018). Yet, we were unable to confidently characterize one of the subroutines of cell death described when analyzing the transcriptional profile of *CLas*^+^ psyllids, once the molecular mechanisms involved were not all represented. However, we can confidently report the activation/inhibition of the expression of genes involved in processes of regulated cell death. The high expression of the lysosomal cathepsin L1 (DN15698_c4_g11_i2) in *A2CLas*^+^ psyllids would support the activation of lysosome-dependent cell death, which requires intracellular perturbations to permeabilize the lysosomal membrane to result in the release of cytosolic cathepsins. Mitochondrial outer membrane permeabilization and caspases are not necessarily required for this process of cell death, and lysosome-dependent cell death is an important response to pathophysiological conditions induced by intracellular pathogens (Galluzzi et al 2018).

The overexpression of the psyllid BCL2/adenovirus E1B 19 kDa protein-interacting protein 3 (BNIP3) (DN_20168_c2_g3_i4) in *A3CLas*^+^ also supports the gut epithelium of *CLas*^+^ psyllids go into lysosome-dependent cell death. BNIP3 are pro-apoptotic proteins that open the pores of mitochondrial outer membrane, resulting in mitochondria dysregulation due the loss of membrane potential and ROS production in mitochondria (Vande Velde et al 2000, Moy & Cherry 2013). But BNIP3 oligomerization with BCL2 has been demonstarted not to be required for cell death (Vande Velde et al. 2000). Besides, the down-regulation of the psyllid senescence-associated protein transcript (DN16041_c1_g1_i2) in the gut of *A3CLas*^+^ psyllids demonstrates CLas suppresses cell senescence and the further dysfunctional growth of the epithelial cells due the activation of senescence-associated secretory phenotype (Münch et al 2008, Ito & Igaki 2016). Cellular senescence is not considered a form of regulated cell death (Galluzzi et al 2018).

The differential expression of a putative juvenile hormone binding protein (JHBP) (DN21423_c1_g12_i2; DN21423_c1_g12_i1) in *CLas*^+^ psyllids (down-regulation in the gut of *A2CLas*^+^; up-regulation in *A3CLas*^+^) demonstrates the epithelial cells count with different levels of hormonal stimulation. JHBPs are proteins that act as shuttles for juvenile hormone (JH) in the hemolymph, avoiding JH degradation by JH-esterases and JH-epoxide hydrolases (Zalewska et al., 2009). JH is produced and released primarily by neurosecretory cells of the *corpora allata* of the central nervous system, and the availability of JHBP in the gut, if its activity in the gut is the same played in the hemolymph, indicates JH is alo available in the gut epithelial cells. JH has recently been shown to be produced by intestinal stem cells and enteroblasts of the gut epithelium of *D. melanogaster,* and to regulate cell growth and survival. The local JH activity was also shown relevant for damage response by gut cells, playing important roles in gut homeostasis (Rahman et al 2017). If the pattern of expression of JHBP correlates with the availability of JH we can argue that at the same time CLas acts on the regulation of the gut epithelium by suppressing cell senescence, it can also induce the proliferation of stem cells in order to replace cells that were damaged due the release of CLas cell into the hemocoel and/or entered the lysosome-mediated cell death as discussed earlier.

The observed up-regulation of the transcriptional levels of nocturnin (DN19927_c10_g1_i6) in the gut of *A2CLas*^+^ psyllids suggests that this circadian rhythm effector protein may be acting together with JH in regulating gut genes, just as the joint action of the circadian genes and JH in the regulation of genes acting on the transitional states of *Pyrrhocoris apterus* adults to reproductive diapause or not (Bajgar et al. 2013). Nocturnin is classified as a deadenylase acting on the catalysis of poly(A) tail of target mRNAs and/or targeting noncoding RNAs, and has been implicated in metabolic regulation, development and differentiation (Hughes et al. 2018). But recent studies with *curled,* the nocturnin ortholog in *Drosophila* proved nocturnin is an NADP(H)2’-phosphatase acting on the conversion of the dinucleotide NADP^*+*^ into NAD^*+*^ and NADPH into NADH, regulating mitochondrial activity and cellular metabolism in response to circadian clock (Estrella et al. 2019). NADP(H)2’- phosphatase activity of nocturnin was later reported in vertebrates, with the demonstration of the colocalization of nocturnin in mitochondria, cytosol and endoplasmic reticulum-bound pools depending on the isoform (Laothamatas et al. 2020). Thus, nocturnin acts as a regulator of the intracellular levels of NADP(H) and the oxidative stress response. The role of nocturnin in the regulation of cell metabolism and the oxidative stress response supports the required energy supply to sustain the increased gene expression activity in *CLas*^+^ gut epithelium cells.

The expression profile of CLas infecting the gut epithelium of *A3CLas*^+^ psyllids demonstrates CLas has an increased protein synthesis activity as expected by the overexpression of DEAD/DEAH box helicase (DN4052_c0_g1_i1), a protein involved in ribosome biogenesis, RNA turnover and translation initiation (Redder et al., 2015). Increased protein activity is also supported by the increased expression of molecular chaperones, such as the trigger factor (DN13945_c0_g1_i1), GroEL (DN14757_c0_g1_i1) and DnaK (DN15220_c0_g1_i1), which prevent protein misfolding and aggregation (Agashe et al 2004, Merz et al. 2006).

### CLas multiprotein complexes

The protein HlyD family efflux transporter periplasmic adaptor subunit (DN11766_c0_g1_i1) and the outer membrane factor translocation protein TolB (DN15087_c0_g1_i3) are essential components for functioning the pump of tripartite efflux systems. HlyD connects primary and secondary inner membrane transporters to the outer membrane factor TolB. CLas expressed several inner membrane transporters belonging to the MSF and ABC families. TolB has been shown to interact with a range of proteins including cell-killing proteins (Carr et al. 2000, Loftus et al. 2006). In *Xylella fastidiosa* TolB was shown to be involved in biofilm development (Santos et al. 2015).

### Additional CLas transcriptional regulators

The detection of several other uncharacterized response regulators (DN10585_c0_g1_i1; DN10585_c0_g2_i1; DN12200_c0_g1_i1; DN58285_c0_g1_i1) and transcriptional regulators (DN11066_c0_g1_i1; DN15145_c0_g1_i2; DN15145_c0_g1_i5) indicates other CLas two-component systems are also being activated by environmental stimuli, such as the two-component system sensor histidine kinase AtoS (DN14305_c0_g2_i1; DN15701_c0_g1_i1). The two-component system sensor histidine kinase AtoS is a member of the AtoS/AtoC regulatory system, but we did not identify the AtoC response regulator in the CLas transcriptome. This regulatory system is better known by the induction of AtoS by acetoacetate. AtoS then phosphorylates AtoC that will in turn stimulate the expression of the atoDAEB operon for the catabolism of short chain fatty acids. We could not reliably identify transcripts belonging to the atoDAEB operon to demonstrate CLas requirements for short chain fatty acids. We propose that the activation of the AtoS/AtoC regulatory system is instead acting on the regulation of the flagellar regulon, regulating the expression of CLas genes involved in cell motility and chemotaxis, as reported for *Escherichia coli* in response to acetoacetate or spermidine (Theodorou et al. 2012).

### Cross-talk on nutrient deficiency

The high expression of sulfonate ABC transporter permease (DN15690_c0_g1_i2) in all *CLas*^+^ psyllids suggests CLas signals the host to supply its requirements for sulfur. Sulfur is a ubiquitous element involved in a number of different processes in organisms (Beinert 2000). The sulfonate ABC transporter permease is involved in sulfur/sulfonate uptake, and the higher expression observed in *A3CLas*^+^ as compared to *A1CLas*^+^ points for an increased demand late in the infection process.

Sulfur is also used in the biogenesis of Fe-S clusters, which are produced in conditions of oxidative stress and iron deprivation. The biogenesis of Fe-S clusters was observed by the expression of two unclassified cysteine sulfurases (DN12917_c0_g1_i1 and DN35541_c0_g1_i1), the cysteine desulfuration protein SufE (DN1144_c0_g1_i1), the Fe-S cluster assembly protein SulfB (DN14252_c0_g1_i1), and the iron-sulfur cluster carrier protein ApbC (DN17762_c4_g2_i3) (Layer et al. 2007, Roche et al. 2013). All four genes had an increased expression in *A3CLas*^+^ as compared to *A1CLas*^+^. Sulfur requirements by CLas could also be used for protection against oxidative stress as demonstrated by the expression of the thioredoxin-dependent thiol peroxidase (DN10526_c0_g1_i1) (Lu & Holmgren 2014; Wang et al. 2020).

The high expression of CLas phosphate ABC transporter permease subunit PstC (DN40256_c0_g1_i1; DN67283_c0_g1_i1), putative two-component sensor histidine kinase transcriptional regulatory protein (DN14305_c0_g2_i1), two-component sensor histidine kinase (PhoR) (DN28075_c0_g1_i1; DN67626_c0_g1_i1), two component response regulator protein (PhoB) (DN14869_c0_g1_i1), alkaline phosphatase (DN12828_c0_g1_i2; DN39297_c0_g1_i2) and NTP pyrophosphohydrolase (DN2128_c0_g1_i1) demonstrates CLas cells are exposed to phosphate restriction while infecting the gut epithelium of adult psyllids. Phosphate is generally the major source of phosphorus to bacteria, a vital nutrient for living organisms. Phosphorus is important in several processes (energy metabolism, intracellular signaling, among others). Misregulation of phosphorus availability in bacteria is sensed by the PhoB/PhoR two-component regulatory system, leading to the activation of the Pho regulon. The Pho regulon regulates the expression of other genes, producing phenotypes that can differ in morphology and virulence in response to phosphate deprivation (Lamarche at al. 2008, Santos-Beneit 2015). Thus, the expression of both members of the PhoB/PhoR two component regulatory system and the phosphate ABC transporter permease subunit PstC, that is in charge to transport extracellular phosphate to the cytosol, proves CLas was exposed to phosphate concentrations below 4 μM, a threshold that generally turns on the expression of the response regulator PhoB. Furthermore, the expression of alkaline phosphatase and NTP pyrophosphohydrolase, which both act on phosphorus-containing substrates (Lamarche et al. 2008) demonstrates CLas increased the catabolism of substrates capable of releasing phosphorus nutrient.

The requirement of CLas for phosphate is also demonstrated by the up-regulation of the psyllid glycerophosphocholine phosphodiesterase GPCPD1 gene, as observed by the increased abundance of its transcript (DN15741_c0_g2_i2) in *A3CLas*^+^. The contribution of GPCPD1 in phosphate production results from the downstream processing of GPCPD1 hydrolysis products of glycerophosphocholine, choline and glycerolphosphate. At the same time glycerophosphocholine hydrolysis can provide phosphates for cell metabolism, the choline can serve as as substrate for phosphatidylcholine synthesis. Phosphatidylcholine is a major lipid component of membranes of eukaryotes, although several bacterial symbionts and pathogens carry phosphatidylcholine synthases, showing their requirement for this lipid component of cell membranes, including those belonging to *Rhizobiaceae* as *Ca*. Liberibacter. Choline serves as substrate for the osmoprotectant glycine betaine synthesis and as an energy substrate to support cell growth in *Rhizobiaceae* (Sohlenkamp et al., 2003; Dupont et al., 2004).

The role of host-produced choline in CLas metabolism is supported by the up-regulation of CLas glycine/betaine ABC transporter substrate-binding protein (DN15178_c0_g1_i1) in the gut of *A3CLas*^+^. ABC transporters were reported to have several important roles in *Ca.* Liberibacter asiaticus, from the importation of nutrients like choline to the exportation of virulence factors (Li et al., 2012). Thus, we believe the increased activity of GPCPD1 in *A3CLas*^+^ is a result of the recycling of glycerophosphocholine for choline utilization in the recovery of membranes of the gut epithelial cells of the host psyllid as a response to cell infection and/or in the supplementation of choline to CLas metabolism.

CLas is known to induce profound physiological changes in citrus plants, affecting the nutritional content of CLas-infected plants to psyllids, which have in turn altered transcriptional profiles and protein abundance when feeding on CLas-infected and health citrus plants (Fu et al. 2016, Ramsey et al. 2017). Therefore, it is impossible to distinguish if the indication of nutritional restriction CLas encounters when infecting the gut epithelium of psyllids is an indirect effect of the host plant on the psyllids or if this is a direct response of psyllids to avoid CLas infection.

## 4. Material and Methods

### 4.1. *Diaphorina citri* rearing and plants

A colony of CLas-free Asian Citrus Psyllid (ACP) was initiated with insects collected from *Murraya paniculata* (L.) Jack, syn. *Murraya exotica* L. (Sapindales: Rutaceae) in the state of São Paulo, Brazil in 2009.

### 4.2. *Diaphorina citri* gut collection and RNA extraction

Adults of *D. citri* (7 to 10 days after emergence) were transferred to uninfected and CLas-infected citrus plants [*Citrus* x *sinensis* (L.) Osbeck, grafted in ‘Rangpur’ lime (*C*. x *limonia* Osbeck)] for an exposure period (EP) of 1, 2, 3, 4, 5 and 6 days. For each exposure period, three biological replicates were collected (1 replicate = 100 individuals). Third-instars of *D. citri* were also transferred to CLas-infected and uninfected (control) citrus plants for collection of RNA and differential transcriptional analysis. Nymphs were allowed to feed for 4 days on CLas-infected and control plants. Afterwards, three biological replicates/treatment (1 replicate = 200 nymphs) were collected and stored in RNALater for further processing and analysis.

After each exposure time, adults were collected and the gut dissected under aseptic conditions. A similar procedure was used for nymphs, but in this case insects we opted by extracting RNA from whole nymphs, as initial attempts to dissect the suitable gut samples for downstream analysis were very time-consuming and had a low success rate.

The obtained guts/nymphs were stored in RNALater (Invitrogen/ThermoFisher Scientific, Waltham, MA, EUA) at −80°C until RNA extraction. Total RNA extraction was performed using the SV RNA Isolation System Kit (Promega), following the manufacturers’ recommendations. Samples were lysed in 175 μL of RNA lysis buffer added with β-mercaptoethanol and 350 μL of RNA dilution buffer for tissue disruption using a TissueLyser II LT^TM^ (Qiagen). Samples were incubated at 70°C for 3 min, centrifuged (12,000 *g* × 10 min × 4°C), trapped in a column and washed with 95% ethanol by centrifugation (12,000 *g* × 10 min × 4°C). Total RNA was recovered in 350 μL of RNA wash solution following centrifugation (12,000 *g* × 1 min × 4°C). Afterwards, samples were treated with 50 μL of DNAse mix (40 μL buffer ‘yellow core’+ 5 μL 0.09 M MnCl_2_ + 5 μL DNAse I) for 15 min at room temperature. Samples were added with 200 μL of DNAse stop solution and subjected to centrifugation (13,000 *g* × 1 min × 4°C). Samples were washed in 600 μL cleaning solution followed by a second wash in 250 μL following centrifugation (12,000 *g* × 1 min × 4°C). The pelleted RNA was recovered in 30 μL of nuclease-free water, and RNA concentration verified using NanoDrop V 3.8.1 (ThermoFischer).

RNA samples containing residual DNA contaminants were further treated with 2 μL of buffer (10×) and 2 μL of Turbo^TM^ DNAse (2 U/μl) (Ambion®). Samples were incubated at 37°C for 30 min followed by DNAse inactivation by adding 2 μL of DNAse Inactivation Reagent (Ambion®). After 5 min at room temperature, samples were centrifuged (13,000 g × 1.5 min × 4°C) and the supernatant collected and stored at −80°C. DNA elimination was confirmed by testing the amplification of the wingless gene (*wg*) of ACP (Manjunath et al., 2008).

All plants and insects were subjected to quantitative PCR analysis for verification of CLas infection using the TaqMan qPCR Master Mix Kit (Ambion®) following Li et al. (2006a).

In insects, the average Ct value found for the *wg* gene in the libraries was 30.7, while the average Ct value in plants for the 16S rRNA gene from CLas was 29.6. Samples with a Ct value under 35 were considered positive for CLas.

After confirmation of the *Ca.* L. asiaticus infection status of each sample, RNA obtained from the gut samples of adults were pooled in equimolar concentrations to yield three exposure periods: *i) A1CLas*^−^: adults that fed on healthy citrus plant (CLas^−^) for 1-2 days; *ii) A2CLas*^−^: for 3-4 days; *iii) A3CLas*^−^: for 5-6 days; *iv) A1CLas*^+^: adults that fed on infected citrus plant (CLas^*+*^) for 1-2 days; *v) A2CLas*^+^: for 3-4 days; and *vi) A3CLas*^+^: for 5-6 days. In the case of the whole nymphs, two samples were produced: *N1CLas*^+^: for nymphs after 4 days of feeding on CLas-infected citrus plants, and *N1CLas*^−^: for nymphs after 4 days of feeding on control plants.

Samples were subjected to mRNA enrichment through eukaryote and prokaryote rRNA removal using the Ribo-Zero rRNA Removal Epidemiology Kit (Illumina®), following the manufacturers’ instructions. RNA integrity was confirmed using the Agilent Bioanalyzer 1000 (Agilent Technologies).

### 4.4. Library preparation and sequencing

cDNA libraries were prepared for sequencing using the cDNA TruSeq RNA Library Prep Kit (Illumina®) following a paired-end (2 × 100 bp) strategy. The cDNA produced was end-repaired and adenosine was added at the 3′ end of each cDNA fragment to guide the ligation of specific adapters. Adapters consisted of primers for transcription and a specific index to code each sample. Samples were enriched with limited-cycle PCR and analyzed to confirm the success of sample preparation before sequencing using the Illumina© HiScanSQ platform available at the Multiusers Center of Agricultural Biotechnology at the Department of Animal Sciences, ESALQ/USP.

### 4.5. *De novo* transcriptome assembly

Reads quality were visualized in FastQC software (Andrews, 2010) before adapters removal and quality filtering by trimming the leading (LEADING:3) and trailing (TRAILING:3) nucleotides until the quality was higher than 3, and then using a sliding window of 4 nucleotides and trimming when scores were lower than 22 (SLIDINGWINDOW 4:22). Quality filtering was done using Trimmomatic-0.36 (Bolger et al., 2014).

All reads obtained were used to assemble a *de novo* transcriptome using the pipeline available in the Trinity-v.2.4.0 software (Haas et al., 2013). Both paired and unpaired trimmed and quality-filtered reads were used to assemble the *de novo* transcriptome, which was further used as the reference transcriptome for the RNA-Seq experiments. Assemblage was obtained using normalization of the reads coverage (<50) and the minimum contig size selected was 200 nucleotides.

The transcripts obtained were functionally annotated using the BlastX algorithm for putative identification of homologous sequences, with an *e*-value cut-off < 10^−3^. Annotated sequences were curated and grouped into categories according to their function using Blast2Go® (Conesa et al., 2005) with an *e*-value cut-off < 10^−6^ and EggNOG-mapper 4.5.1 (Huerta-Cepas et al., 2017). Transcripts putatively identified as belonging to insects and *Ca.* L. asiaticus were checked against the KEGG database (Kyoto Encyclopedia of Genes and Genomes) (Kanehisa & Goto, 2000) to verify the metabolic pathways represented in the obtained *de novo* transcriptome. Transcripts of *Diaphorina citri*, *Ca.* Liberibacter spp. and *Wolbachia* spp. were filtered using Blast2go version Pro (Götz et al. 2008).

### 4.6. Differential gene expression analyses

Changes in the pattern of gene expression were evaluated separately in gut of ACP adults infected or not by *Ca*. L. asiaticus using the CLC Genomics Workbench 20.0 software (QIAGEN, Aarhus, Denmark). Reads from each library were counted against the *de novo* transcriptome, and counts were normalized as transcripts per million reads (TPM). TPM values for each sample were used to calculate fold-change ratios for comparative analyses of the gene expression of control (CLas^−^) versus CLas-infected insects (CLas^+^) within each feeding interval (1-2 d, 3-4 d, 5-6 d). Only the transcript that were counted in two of the replicates of a particular treatment were further taken for comparative analysis. Data were analyzed using multifactorial statistics based on a negative binomial Generalized Linear Model (GLM). The values of fold change obtained were corrected with False Discovery Rate (FDR) method and only transcripts that showed *p*-value ≤0.05 and log fold change > |2| between treatments were considered differentially expressed.

## Acknowledgements

We are grateful to FAPESP for providing a post-doctoral fellowship to FMMB (grant: 2018/24234-4) and a research grant to FLC, NAW and LP to support this research (grant: 15/07011-3). To CNPq for the PhD fellowship to JCD (grant: 141042/2017-6). Authors also acknowledge the Fundecitrus support personnel team that provided the required assistance for conducting this research.

## Author’s contribution

FLC and NAW designed the experiments; FLC, NAW and LP secured the funds; JCD collected insect samples and extracted RNA; BLM processed the sequencing data and assembled the *de novo* transcriptome; FMMB analyzed the data; FMMB and FLC wrote the paper; all authors commented the initial draft and approved the final version of this manuscript.

---

**S1 Table.**
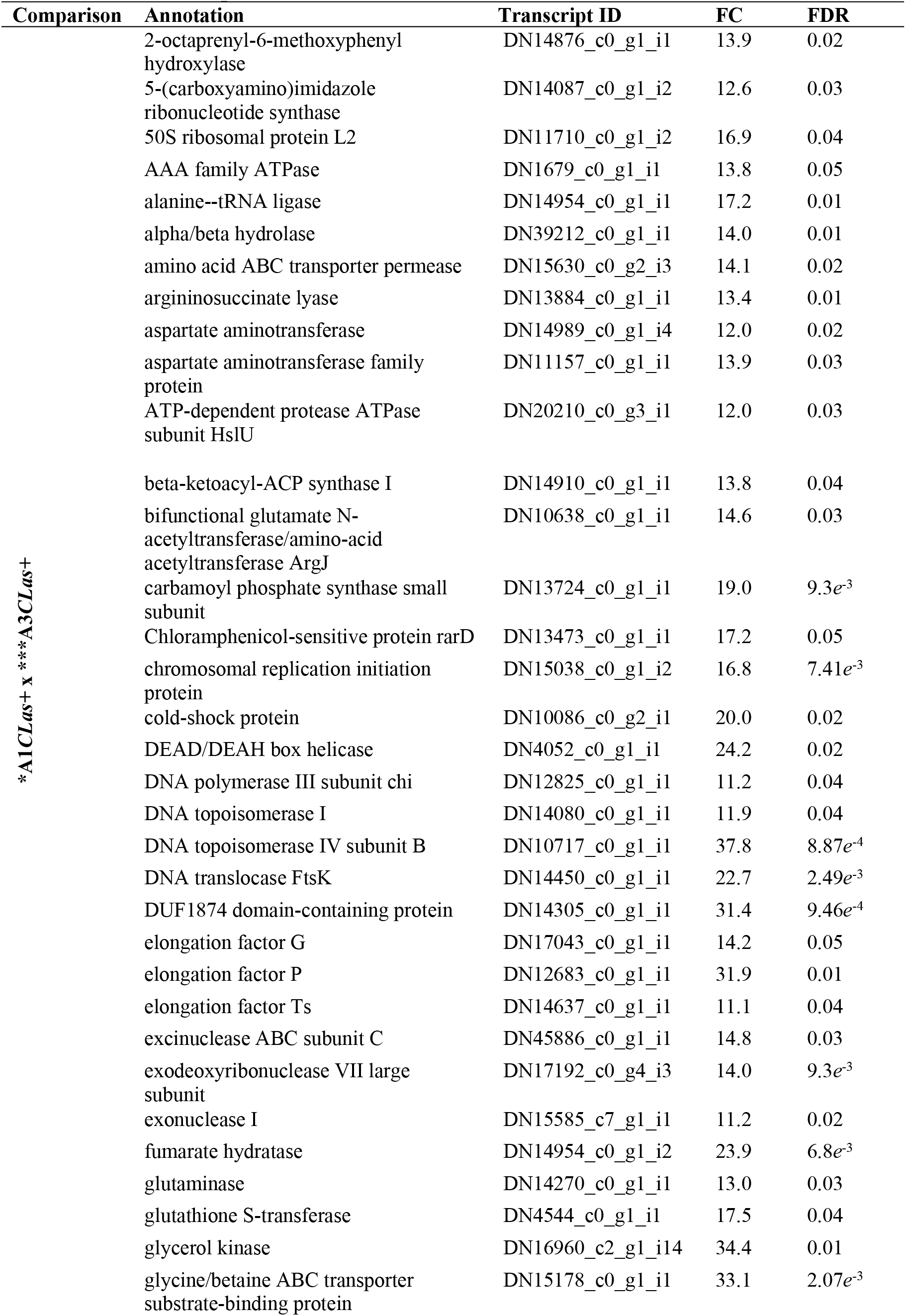
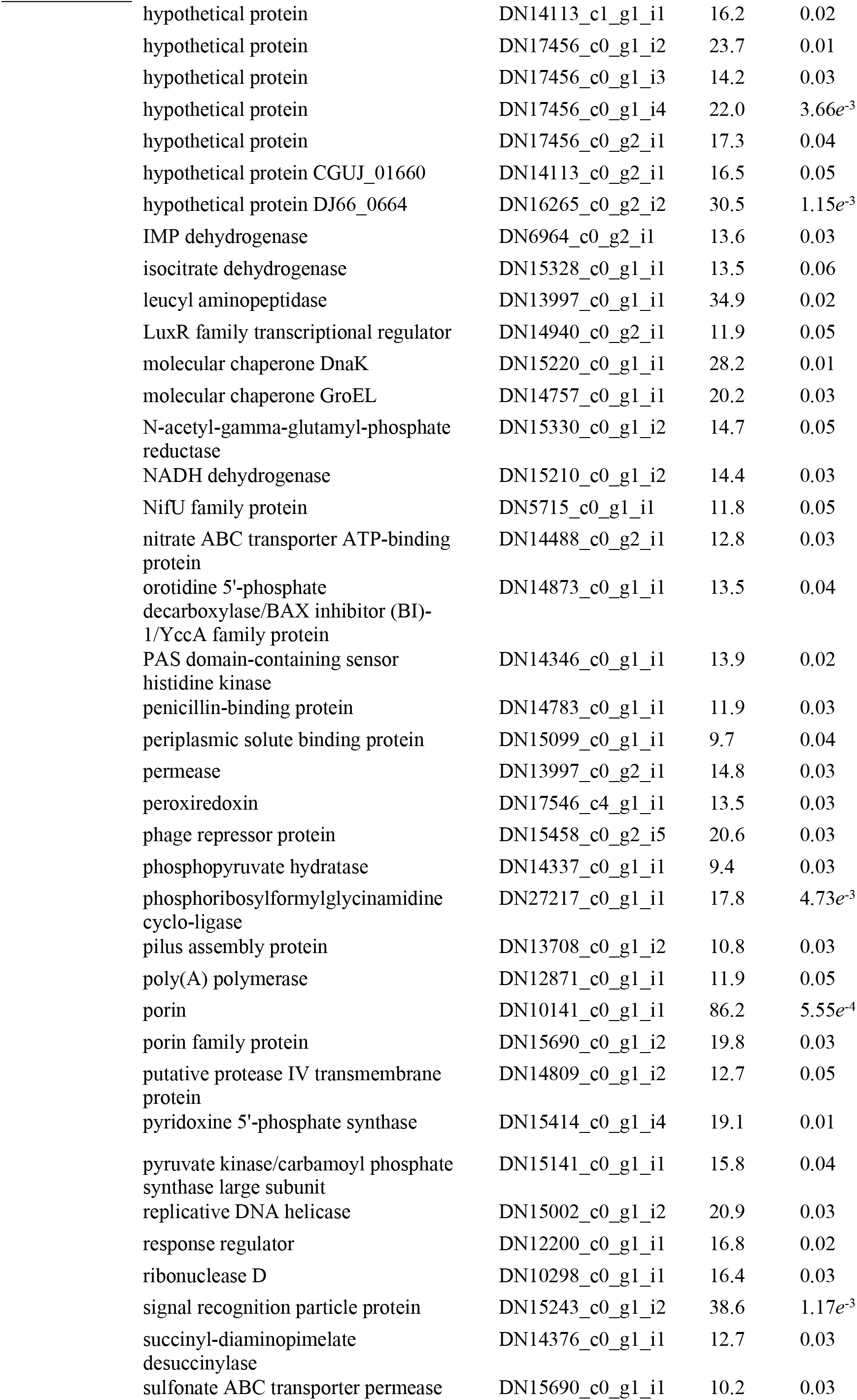
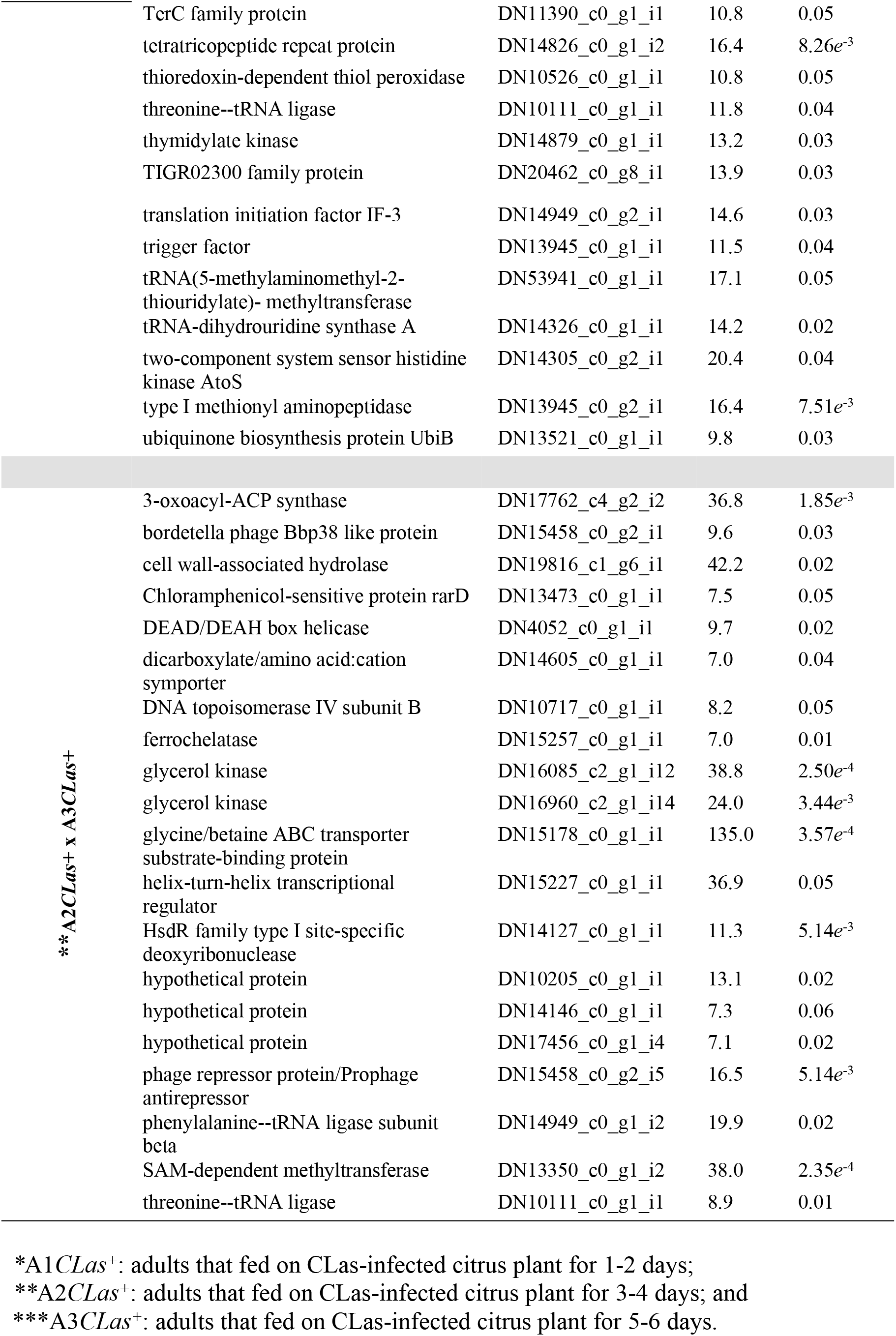
Differentially expressed CLas genes in the gut of *Diaphorina citri* adults after 1-2 d (*A1CLas*^+^), 3-4 d (*A2CLas*^+^) and 5-6 d (*A3CLas*^+^) of infection by feeding on CLas-infected citrus plants.

**Fig S1.**
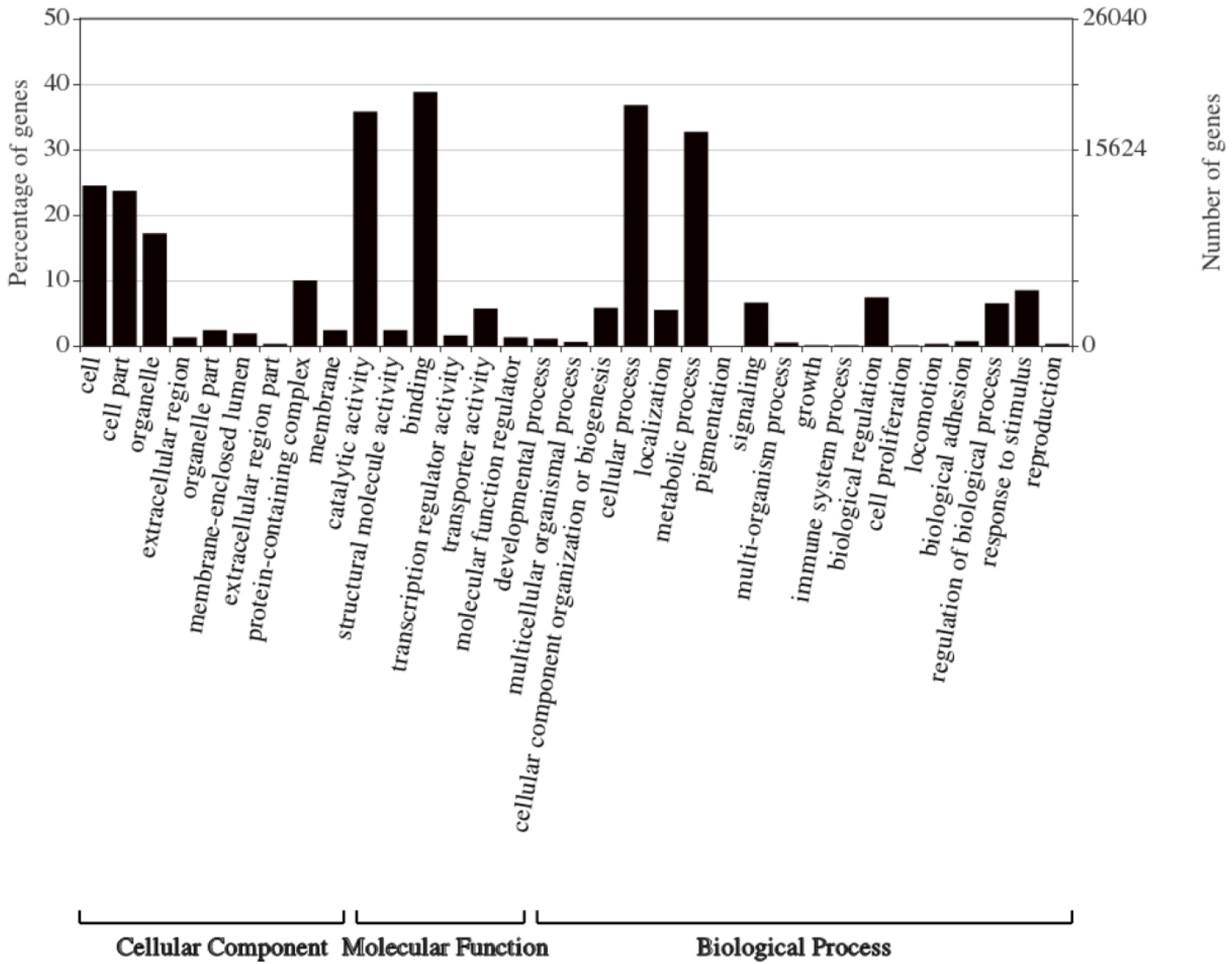
The analysis of the 248,850 contigs was conducted by Blast2Go, using Blastx to recover annotations with significant homology from the NCBI. All terms ‘Biological Processes’, ‘Molecular Function’ and ‘Cellular Component’ at level 2 are represented as percent over total number of sequences.

**Fig S2.**
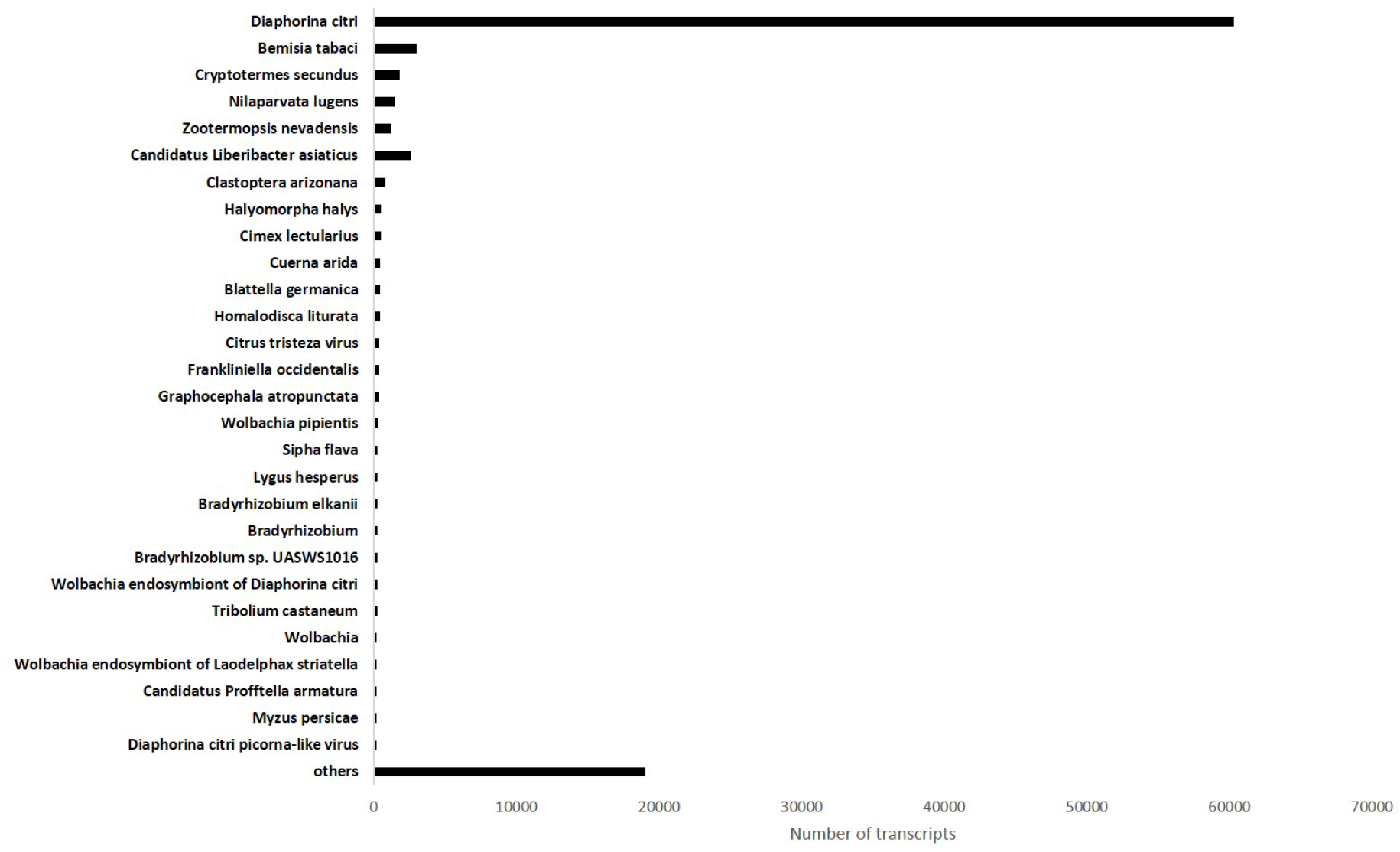
Distribution of alignments and transcripts of *de novo* assembly of *Diaphorina citri* gut that fed on health and CLas-infected citrus plant, after search for similarity in NCBI databank.

## References

Abebe E, Gugsa G, Ahmed M. Review on Major Food-Borne Zoonotic Bacterial Pathogens. J Trop Med. 2020;2020:4674235. doi: 10.1155/2020/4674235.

Agashe VR, Guha S, Chang HC, Genevaux P, Hayer-Hartl M, Stemp M, et al. Function of trigger factor and DnaK in multidomain protein folding: increase in yield at the expense of folding speed. Cell. 2004;117(2):199–209. doi: 10.1016/s0092-8674(04)00299-5.

Ajene IJ, Khamis FM, van Asch B, Pietersen G, Seid N, Rwomushana I, et al. Distribution of *Candidatus* Liberibacter species in Eastern Africa, and the First Report of *Candidatus* Liberibacter asiaticus in Kenya. Sci Rep. 2020;10:3919. doi: 10.1038/s41598-020-60712-0.

Amherd R, Hintermann E, Walz D, Affolter M, Meyer UA. Purification, cloning, and characterization of a second arylalkylamine N-acetyltransferase from *Drosophila melanogaster*. DNA Cell Biol. 2000;19(11):697–705. doi: 10.1089/10445490050199081.

Ammar E, Shatters Jr RG, Lynch C, Hall DG. Detection and Relative Titer of *Candidatus* Liberibacter asiaticus in the salivary glands and alimentary canal of *Diaphorina citri* (Hemiptera: Psyllidae) vector of Citrus Huanglongbing Disease. Ann Entomol Soc Am. 2011b;104(3):526–533. doi: 10.1603/AN10134.

Ammar E-D, Achor D, Levy A. Immuno-ultrastructural localization and putative multiplication sites of Huanglongbing bacterium in Asian Citrus Psyllid *Diaphorina citri*. Insects. 2019;10(12):422. doi: 10.3390/insects10120422.

Ammar E-D, George J, Sturgeon K, Stelinski LL, Shatters RG. Asian citrus psyllid adults inoculate Huanglongbing bacterium more efficiently than nymphs when this bacterium is acquired by early instar nymphs. Sci Rep. 2020;10(1)18244. doi: 10.1038/s41598-020-75249-5.

Ammar E-D, Hall DG, Shatters RG Jr. Ultrastructure of the salivary glands, alimentary canal and bacteria-like organisms in the Asian citrus psyllid, vector of citrus Huanglongbing disease bacteria. J Microsc Ultrastruct. 2017;5:9–20. doi: 10.1016/j.jmau.2016.01.005.

Ammar E-D, Ramos JE, Hall DG, Dawson WO, Shatters RG Jr. Acquisition, replication and inoculation of *Candidatus* Liberibacter asiaticus following various acquisition periods on Huanglongbing-Infected citrus by nymphs and adults of the Asian Citrus Psyllid. PLoS One. 2016;11(7): e0159594. doi: 10.1371/journal.pone.0159594.

Ammar E-D, Shatters Jr RG, Hall DG. Localization of *Candidatus* Liberibacter asiaticus, associated with Citrus Huanglongbing Disease, in its psyllid vector using fluorescence in situ hybridization. J Phytopathol. 2011a;159:726–734. doi: 10.1111/j.1439-0434.2011.01836.x.

Anderson KV. Toll signaling pathways in the innate immune response. Curr Opin Immunol. 2000;12(1):13–19. doi: 10.1016/s0952-7915(99)00045-x.

Andrade M, Li JY, Wang N. *Candidatus* Liberibacter asiaticus: virulence traits and control strategies. Trop Plant Pathol. 2020b;45:285–297. doi: 10.1007/s40858-020-00341-0.

Andrade M, Wang N. (2019) The Tad Pilus apparatus of ‘*Candidatus* Liberibacter asiaticus’ and its regulation by VisNR. Mol Plant Microbe Interact. 2019;32(9):1175–1187. doi: 10.1094/MPMI-02-19-0052-R.

Andrade MO, Pang Z, Achor DS, Wang H, Yao T, Singer BH, Wang N. The flagella of ‘*Candidatus* Liberibacter asiaticus’ and its movement *in planta*. Mol Plant Pathol. 2020a;21:109–123. doi: 10.1111/mpp.12884.

Andrews S. FastQC: a quality control tool for high throughput sequence data [software]. 2010 [cited 2021 Feb 26]. Available from: http://www.bioinformatics.babraham.ac.uk/projects/fastqc.

Bajgar A, Jindra M, Dolezel D. Autonomous regulation of the insect gut by circadian genes acting downstream of juvenile hormone signaling. Proc Natl Acad Sci U S A. 2013;110(11):4416–4421. doi: 10.1073/pnas.1217060110.

Bardy SL, Ng SYM, Jarrell KF. Prokaryotic motility structures. Microbiology. 2003;149:295–304. doi: 10.1099/mic.0.25948-0.

Bassanezi RB, Lopes AS, Miranda MP, Wulff NA, Volpe HXL, Ayres, AJ. Overview of citrus huanglongbing spread and management strategies in Brazil. Trop Plant Pathol. 2020;45:251–264. doi: 10.1007/s40858-020-00343-y.

Baumann P. Biology bacteriocyte-associated endosymbionts of plant sap-sucking insects. Annu Rev Microbiol. 2005;59:155–89. doi: 10.1146/annurev.micro.59.030804.121041.

Beinert H. A tribute to sulfur. Eur J Biochem. 2000;267:5657–5664. doi: 10.1046/j.1432-1327.2000.01637.x.

Bendix C, Lewis JD. The enemy within: phloem-limited pathogens. Mol Plant Pathol. 2018 Jan;19(1):238–254. doi: 10.1111/mpp.12526.

Benguettat O, Jneid R, Soltys J, Loudhaief R, Brun-Barale A, Osman D, Gallet A. The DH31/CGRP enteroendocrine peptide triggers intestinal contractions favoring the elimination of opportunistic bacteria. PLoS Pathog. 2018;14(9):e1007279. doi: 10.1371/journal.ppat.1007279.

Bhattacharyya S, Feferman L, Tobacman JK. Arylsulfatase B regulates versican expression by galectin-3 and AP-1 mediated transcriptional effects. Oncogene. 2014;33(47):5467–5476. doi: 10.1038/onc.2013.483.

Bolger AM, Lohse M, Usadel B. Trimmomatic: a flexible trimmer for Illumina sequence data. Bioinformatics. 2014;30(15):2114–2120. doi: 10.1093/bioinformatics/btu170.

Bové JM, Garnier M. Phloem-and xylem-restricted plant pathogenic bacteria. Plant Sci. 2003;164:423–438. doi: 10.1016/S0168-9452(03)00033-5.

Bové JM. Huanglongbing: A destructive newly emerging, century-old disease of citrus. J Plant Pathol. 2006;88:7–37. doi: 10.4454/jpp.v88i1.828.

Brodbeck D, Amherd R, Callaerts P, Hintermann E, Meyer UA, Affolter M. Molecular and biochemical characterization of the aaNAT1 (Dat) locus in *Drosophila melanogaster*: differential expression of two gene products. DNA Cell Biol. 1998;17(7):621–633. doi: 10.1089/dna.1998.17.621.

Brunner E, Peter O, Schweizer L, Basler K. *pangolin* encodes a Lef-1 homologue that acts downstream of Armadillo to transduce the Wingless signal in *Drosophila*. Nature. 1997;385(6619):829–833. doi: 10.1038/385829a0.

Buchon N, Broderick NA, Lemaitre B. Gut homeostasis in a microbial world: insights from *Drosophila melanogaster*. Nat Rev Microbiol. 2013;11(9):615–26. doi: 10.1038/nrmicro3074.

CABI/EPPO. *Candidatus* Liberibacter asiaticus [Distribution map]. 4th ed. Wallingford: CABI.; 2017.

Caccia S, Casartelli M, Tettamanti G. The amazing complexity of insect midgut cells: types, peculiarities, and functions. Cell Tissue Res. 2019;377(3):505–525. doi: 10.1007/s00441-019-03076-w.

Carr S, Penfold CN, Bamfold V, James R, Hemmings AM. The structure of TolB, an essential component of the tol-dependent translocation system, and its protein–protein interaction with the translocation domain of colicin E9. Structure. 2000;8:57–66. doi: 10.1016/S0969-2126(00)00079-4.

Celli J, Gorvel JP. Organelle robbery: *Brucella* interactions with the endoplasmic reticulum. Curr Opin Microbiol. 2004;7(1):93–97. doi: 10.1016/j.mib.2003.11.001.

Cerveny L, Straskova A, Dankova V, Hartlova A, Ceckova M, Staud F, Stulik J. Tetratricopeptide repeat motifs in the world of bacterial pathogens: role in virulence mechanisms. Infect Immun. 2013;81(3):629–635. doi: 10.1128/IAI.01035-12.

Cicero JM, Fisher TW, Qureshi JA, Stansly PA, Brown JK. Colonization and Intrusive Invasion of Potato Psyllid by ‘*Candidatus* Liberibacter solanacearum’. Phytopathology. 2017;107(1):36–49. doi: 10.1094/PHYTO-03-16-0149-R.

Conesa A, Götz S, García-Gómez JM, Terol J, Talón M, Robles M. Blast2GO: a universal tool for annotation, visualization and analysis in functional genomics research. Bioinformatics. 2005;21(18):3674–3676. doi: 10.1093/bioinformatics/bti610.

Cordero JB, Stefanatos RK, Scopelliti A, Vidal M, Sansom OJ. Inducible progenitor-derived Wingless regulates adult midgut regeneration in *Drosophila*. EMBO J. 2012;31(19):3901–3917. doi: 10.1038/emboj.2012.248.

Dala-Paula BM, Raithore S, Manthey JA, Baldwin EA, Zhao W et al. Active taste compounds in juice from oranges symptomatic for Huanglongbing (HLB) citrus greening disease. LWT – Food Science Technology. 2018;9:518–525. doi: 10.1016/j.lwt.2018.01.083.

Dala-Paula BM, Plotto A, Bai J, Manthey JA, Baldwin EA, Ferrarezi RS, Gloria MBA. Effect of Huanglongbing or Greening Disease on Orange Juice Quality, a Review. Front Plant Sci. 2019;9:1976. doi: 10.3389/fpls.2018.01976.

Dohrman A, Miyata S, Gallup M, Li JD, Chapelin C, Coste A, et al. Mucin gene (MUC 2 and MUC 5AC) upregulation by Gram-positive and Gram-negative bacteria. Biochim Biophys Acta. 1998;1406(3):251–259. doi: 10.1016/s0925-4439(98)00010-6.

Duan Q, Zhou M, Zhu L, Zhu G. Flagella and bacterial pathogenicity. J Basic Microbiol. 2013;53(1):1–8. doi: 10.1002/jobm.201100335.

Dupont L, Garcia I, Poggi MC, Alloing G, Mandon K, Le Rudulier D. The *Sinorhizobium meliloti* ABC transporter Cho is highly specific for choline and expressed in bacteroids from *Medicago sativa* nodules. J Bacteriol. 2004;186(18):5988–96. doi: 10.1128/JB.186.18.5988-5996.2004.

Erlandson MA, Toprak U, Hegedus DD. Role of the peritrophic matrix in insect-pathogen interactions. J Insect Physiol. 2019;117:103894. doi: 10.1016/j.jinsphys.2019.103894.

Eshwar AK, Guldimann C, Oevermann A, Tasara T. Cold-Shock Domain Family Proteins (Csps) Are Involved in Regulation of Virulence, Cellular Aggregation, and Flagella-Based Motility in *Listeria* monocytogenes. Front Cell Infect Microbiol. 2017;7:453. doi: 10.3389/fcimb.2017.00453.

Estrella MA, Du J, Chen L, Rath S, Prangley E, Chitrakar A, Aoki T, Schedl P, Rabinowitz J, Korennykh A. The metabolites NADP+ and NADPH are the targets of the circadian protein Nocturnin (Curled). Nat Commun. 2019;10(1):2367. doi: 10.1038/s41467-019-10125-z.

Faggioni R, Feingold KR, Grunfeld C. Leptin regulation of the immune response and the immunodeficiency of malnutrition. FASEB J. 2001;15(14):2565–71. doi: 10.1096/fj.01-0431rev.

Fauci AS. Infectious diseases: considerations for the 21st century. Clin Infect Dis. 2001;32(5):675–85. doi: 10.1086/319235.

Fong JNC, Yildiz FH. Biofilm Matrix Proteins. Microbiol Spectr. 2015;3(2):10.1128/microbiolspec.MB-0004-2014. doi: 10.1128/microbiolspec.MB-0004-2014.

Franz A, Shlyueva D, Brunner E, Stark A, Basler K. Probing the canonicity of the Wnt/Wingless signaling pathway. PLoS Genet. 2017;13(4):e1006700. doi: 10.1371/journal.pgen.1006700.

Friedman JR, Dibenedetto JR, West M, Rowland AA, Voeltz GK. Endoplasmic reticulum-endosome contact increases as endosomes traffic and mature. Mol Biol Cell. 2013;24(7):1030–1040. doi: 10.1091/mbc.E12-10-0733.

Fu S, Shao J, Zhou C, Hartung JS. Transcriptome analysis of sweet orange trees infected with *‘Candidatus* Liberibacter asiaticus’ and two strains of Citrus Tristeza Virus. BMC Genomics. 2016;17:349. doi: 10.1186/s12864-016-2663-9.

Galganski L, Urbanek MO, Krzyzosiak WJ. Nuclear speckles: molecular organization, biological function and role in disease. Nucleic Acids Res. 2017;45(18):10350–10368. doi: 10.1093/nar/gkx759.

Galluzzi L, Vitale I, Aaronson SA, Abrams JM, Adam D, Agostinis P, Alnemri ES, et al. Molecular mechanisms of cell death: recommendations of the Nomenclature Committee on Cell Death 2018. Cell Death Differ. 2018;25(3):486–541. doi: 10.1038/s41418-017-0012-4.

García B, Merayo-Lloves J, Martin C, Alcalde I, Quirós LM, Vazquez F. Surface Proteoglycans as Mediators in Bacterial Pathogens Infections. Front Microbiol. 2016;7:220. doi: 10.3389/fmicb.2016.00220.

Ghanim M, Achor D, Ghosh S, Kontsedalov S, Lebedev G, Levy A. *‘Candidatus* Liberibacter asiaticus’ accumulates inside endoplasmic reticulum associated vacuoles in the gut cells of *Diaphorina citri*. Sci Rep. 2017;7(1):16945. doi: 10.1038/s41598-017-16095-w.

Giltner CL, Nguyen Y, Burrows LL. Type IV pilin proteins: versatile molecular modules. Microbiol Mol Biol Rev. 2012;76(4):740–72. doi: 10.1128/MMBR.00035-12.

Girard C, Will CL, Peng J, Makarov EM, Kastner B, Lemm I, et al. Post-transcriptional spliceosomes are retained in nuclear speckles until splicing completion. Nat Commun. 2012;3:994. doi: 10.1038/ncomms1998.

Götz S, García-Gómez JM, Terol J, Williams TD, Nagaraj SH, Nueda MJ, et al. High-throughput functional annotation and data mining with the Blast2GO suite. Nucleic Acids Res. 2008;36(10):3420–35. doi: 10.1093/nar/gkn176.

Graça JV, Douhan GW, Halbert SE, Keremane ML, Lee RF, Vidalakis G, Zhao H. Huanglongbing: An overview of a complex pathosystem ravaging the world’s citrus. J Integr Plant Biol. 2016;58(4):373–87. doi: 10.1111/jipb.12437.

Granger E, McNee G, Allan V, Woodman P. The role of the cytoskeleton and molecular motors in endosomal dynamics. Semin Cell Dev Biol. 2014;31(100):20–9. doi: 10.1016/j.semcdb.2014.04.011.

Gray S, Cilia M, Ghanim M. Circulative, “nonpropagative” virus transmission: an orchestra of virus-, insect-, and plant-derived instruments. Adv Virus Res. 2014;89:141–199. doi: 10.1016/B978-0-12-800172-1.00004-5.

Green ER, Mecsas J. Bacterial Secretion Systems: An Overview. Microbiol Spectr. 2016;4(1):10.1128/microbiolspec.VMBF-0012-2015. doi: 10.1128/microbiolspec.VMBF-0012-2015.

Guo X, Roberts MR, Becker SM, Podd B, Zhang Y, Chua SC Jr, et al. Leptin signaling in intestinal epithelium mediates resistance to enteric infection by *Entamoeba histolytica*. Mucosal Immunol. 2011;4(3):294–303. doi: 10.1038/mi.2010.76.

Haas BJ, Papanicolaou A, Yassour M, Grabherr M, Blood PD, Bowden J, et al. *De novo* transcript sequence reconstruction from RNA-seq using the Trinity platform for reference generation and analysis. Nat Protoc. 2013;8(8):1494–512. doi: 10.1038/nprot.2013.084.

Hammer TJ, Janzen DH, Hallwachs W, Jaffe SP, Fierer N. Caterpillars lack a resident gut microbiome. Proc Natl Acad Sci U S A. 2017;114(36):9641–9646. doi: 10.1073/pnas.1707186114.

Hansson GC. Role of mucus layers in gut infection and inflammation. Curr Opin Microbiol. 2012;15(1):57–62. doi: 10.1016/j.mib.2011.11.002.

Helbig JH, König B, Knospe H, Bubert B, Yu C, Lück CP, et al. The PPIase active site of *Legionella pneumophila* Mip protein is involved in the infection of eukaryotic host cells. Biol Chem. 2003;384(1):125–37. doi: 10.1515/BC.2003.013. PMID: 12674506.

Hijaz F, Al-Rimawi F, Manthey JA, Killiny N. Phenolics, flavonoids and antioxidant capacities in Citrus species with different degree of tolerance to Huanglongbing. Plant Signal Behav. 2020 May 3;15(5):1752447. doi: 10.1080/15592324.2020.1752447.

Hijaz FM, Manthey JA, Folimonova SY, Davis CL, Jones SE, Reyes-De-Corcuera JI. An HPLC-MS characterization of the changes in sweet orange leaf metabolite profile following infection by the bacterial pathogen *Candidatus* Liberibacter asiaticus. PLoS One. 2013;8(11):e79485. doi: 10.1371/journal.pone.0079485.

Hiragaki S, Suzuki T, Mohamed AA, Takeda M. Structures and functions of insect arylalkylamine N-acetyltransferase (iaaNAT); a key enzyme for physiological and behavioral switch in arthropods. Front Physiol. 2015;6:113. doi: 10.3389/fphys.2015.00113.

Hirokawa N. Kinesin and dynein superfamily proteins and the mechanism of organelle transport. Science. 1998;279(5350):519–526. doi: 10.1126/science.279.5350.519.

Holzbaur EL, Vallee RB. DYNEINS: molecular structure and cellular function. Annu Rev Cell Biol. 1994;10:339–372. doi: 10.1146/annurev.cb.10.110194.002011.

Hopkins AM, Walsh SV, Verkade P, Boquet P, Nusrat A. Constitutive activation of Rho proteins by CNF-1 influences tight junction structure and epithelial barrier function. J Cell Sci. 2003;116:725–742. doi: 10.1242/jcs.00300.

Huang W, Reyes-Caldas P, Mann M, Seifbarghi S, Kahn A, Almeida RPP, et al. Bacterial Vector-Borne Plant Diseases: Unanswered Questions and Future Directions. Mol Plant. 2020;13(10):1379–1393. doi: 10.1016/j.molp.2020.08.010.

Huerta-Cepas J, Forslund K, Coelho LP, Szklarczyk D, Jensen LJ, von Mering C, Bork P. Fast Genome-Wide Functional Annotation through Orthology Assignment by eggNOG-Mapper. Mol Biol Evol. 2017;34(8):2115–2122. doi: 10.1093/molbev/msx148.

Hughes KL, Abshire ET, Goldstrohm AC. Regulatory roles of vertebrate Nocturnin: insights and remaining mysteries. RNA Biol. 2018;15(10):1255–1267. doi: 10.1080/15476286.2018.1526541.

Ito T, Igaki T. Dissecting cellular senescence and SASP in Drosophila. Inflamm Regen. 2016;36:25. doi: 10.1186/s41232-016-0031-4.

Jagoueix S, Bove JM, Garnier M. The phloem-limited bacterium of greening disease of citrus is a member of the alpha subdivision of the Proteobacteria. Int J Syst Bacteriol. 1994;44(3):379–786. doi: 10.1099/00207713-44-3-379.

Jing T, Wang F, Qi F, Wang Z. Insect anal droplets contain diverse proteins related to gut homeostasis. BMC Genomics. 2018;19(1):784. doi: 10.1186/s12864-018-5182-z.

Kanehisa M, Goto S. KEGG: kyoto encyclopedia of genes and genomes. Nucleic Acids Res. 2000;28(1):27–30. doi: 10.1093/nar/28.1.27.

Kazmierczak BI, Schniederberend M, Jain R. Cross-regulation of *Pseudomonas* motility systems: the intimate relationship between flagella, pili and virulence. Curr Opin Microbiol. 2015;28:78–82. doi: 10.1016/j.mib.2015.07.017.

Kiefl J, Kohlenberg B, Hartmann A, Obst K, Paetz S, Krammer G, Trautzsch S. Investigation on key molecules of Huanglongbing (HLB)-induced orange juice off-flavor. J Agric Food Chem. 2018;66(10):2370–2377. doi: 10.1021/acs.jafc.7b00892.

Klein DC. Arylalkylamine N-acetyltransferase: “the Timezyme”. J Biol Chem. 2007;282(7):4233–4237. doi: 10.1074/jbc.R600036200.

Knoops B, Argyropoulou V, Becker S, Ferté L, Kuznetsova O. Multiple Roles of Peroxiredoxins in Inflammation. Mol Cells. 2016;39(1):60–4. doi: 10.14348/molcells.2016.2341.

Kocks C. Intracellular motility. Profilin puts pathogens on the actin drive. Curr Biol. 1994;4(5):465–8. doi: 10.1016/s0960-9822(00)00105-6.

Kodrík D, Bednářová A, Zemanová M, Krishnan N. Hormonal regulation of response to oxidative stress in insects-an update. Int J Mol Sci. 2015;16(10):25788–816. doi: 10.3390/ijms161025788.

Kostygov AY, Frolov AO, Malysheva MN, Ganyukova AI, Chistyakova LV, Tashyreva D, et al. *Vickermania* gen. nov., trypanosomatids that use two joined flagella to resist midgut peristaltic flow within the fly host. BMC Biol. 2020;18(1):187. doi: 10.1186/s12915-020-00916-y.

Kruse A, Fattah-Hosseini S, Saha S, Johnson R, Warwick E, Sturgeon K, et al. Combining ’omics and microscopy to visualize interactions between the Asian citrus psyllid vector and the Huanglongbing pathogen *Candidatus* Liberibacter asiaticus in the insect gut. PLoS One. 2017;12(6):e0179531. doi: 10.1371/journal.pone.0179531.

Kuraishi T, Binggeli O, Opota O, Buchon N, Lemaitre B. Genetic evidence for a protective role of the peritrophic matrix against intestinal bacterial infection in *Drosophila melanogaster*. Proc Natl Acad Sci U S A. 2011;108(38):15966–71. doi: 10.1073/pnas.1105994108.

Lakhssassi N, Liu S, Bekal S, Zhou Z, Colantonio V, Lambert K, et al. Characterization of the Soluble NSF Attachment Protein gene family identifies two members involved in additive resistance to a plant pathogen. Sci Rep. 2017;7:45226. doi: 10.1038/srep45226.

Lamarche MG, Wanner BL, Crépin S, Harel J. The phosphate regulon and bacterial virulence: a regulatory network connecting phosphate homeostasis and pathogenesis. FEMS Microbiol Rev. 2008;32(3):461–73. doi: 10.1111/j.1574-6976.2008.00101.x.

Laothamatas I, Gao P, Wickramaratne A, Quintanilla CG, Dino A, Khan CA, Liou J, Green CB. Spatiotemporal regulation of NADP(H) phosphatase Nocturnin and its role in oxidative stress response. Proc Natl Acad Sci U S A. 2020;117(2):993–999. doi: 10.1073/pnas.1913712117.

Layer G, Gaddam SA, Ayala-Castro CN, Ollagnier-de Choudens S, Lascoux D, Fontecave M, Outten FW. SufE transfers sulfur from SufS to SufB for iron-sulfur cluster assembly. J Biol Chem. 2007;282(18):13342–50. doi: 10.1074/jbc.M608555200.

Lehane MJ, Billingsley PF. Biology of the Insect Midgut. Dordrecht: Springer Netherlands; 1996. doi : 10.1007/978-94-009-1519-0.

Li W, Hartung JS, Levy L. Quantitative real-time PCR for detection and identification of *Candidatus* Liberibacter species associated with citrus huanglongbing. J Microbiol Methods. 2006a;66(1):104–15. doi: 10.1016/j.mimet.2005.10.018.

Li Y, Park JS, Deng JH, Bai Y. Cytochrome c oxidase subunit IV is essential for assembly and respiratory function of the enzyme complex. J Bioenerg Biomembr. 2006b;38(5-6):283–91. doi: 10.1007/s10863-006-9052-z.

Li W, Cong Q, Pei J, Kinch LN, Grishin NV. The ABC transporters in *Candidatus* Liberibacter asiaticus. Proteins. 2012;80(11):2614–28. doi: 10.1002/prot.24147.

Liu R, Ochman H. Origins of flagellar gene operons and secondary flagellar systems. J Bacteriol. 2007;189(19):7098–7104. doi: 10.1128/JB.00643-07.

Loftus SR, Walker D, Maté MJ, Bonsor DA, James R, Moore GR, Kleanthous C. Competitive recruitment of the periplasmic translocation portal TolB by a natively disordered domain of colicin E9. Proc Natl Acad Sci U S A. 2006;103(33):12353–8. doi: 10.1073/pnas.0603433103.

Lu H, Zhu J, Yu J, Chen X, Kang L, Cui F. A Symbiotic Virus Facilitates Aphid Adaptation to Host Plants by Suppressing Jasmonic Acid Responses. Mol Plant Microbe Interact. 2020;33(1):55–65. doi: 10.1094/MPMI-01-19-0016-R.

Lu J, Holmgren A. The thioredoxin antioxidant system. Free Radic Biol Med. 2014;66:75–87. doi: 10.1016/j.freeradbiomed.2013.07.036.

Lu ZJ, Huang YL, Yu HZ, Li NY, Xie YX, Zhang Q, et al. Silencing of the Chitin Synthase Gene Is Lethal to the Asian Citrus Psyllid, *Diaphorina citri*. Int J Mol Sci. 2019a;20(15):3734. doi: 10.3390/ijms20153734.

Lu ZJ, Zhou CH, Yu HZ, Huang YL, Liu YX, Xie YX, et al. Potential roles of insect Tropomyosin1-X1 isoform in the process of *Candidatus* Liberibacter asiaticus infection of *Diaphorina citri*. J Insect Physiol. 2019b;114:125–135. doi: 10.1016/j.jinsphys.2019.02.012.

Mackey-Lawrence NM, Petri WA Jr. Leptin and mucosal immunity. Mucosal Immunol. 2012;5(5):472–9. doi: 10.1038/mi.2012.40.

Macnab RM. How bacteria assemble flagella. Annu Rev Microbiol. 2003;57:77–100. doi: 10.1146/annurev.micro.57.030502.090832.

Manjunath KL, Halbert SE, Ramadugu C, Webb S, Lee RF. Detection of *‘Candidatus* Liberibacter asiaticus’ in *Diaphorina citri* and its importance in the management of citrus huanglongbing in Florida. Phytopathology. 2008;98(4):387–396. doi: 10.1094/PHYTO-98-4-0387.

Mansfield J, Genin S, Magori S, Citovsky V, Sriariyanum M, Ronald P, et al. Top 10 plant pathogenic bacteria in molecular plant pathology. Mol Plant Pathol. 2012;13(6):614–29. doi: 10.1111/j.1364-3703.2012.00804.x.

Massenti R, Lo Bianco R, Sandhu AK, Gu L, Sims C. Huanglongbing modifies quality components and flavonoid content of ‘Valencia’ oranges. J Sci Food Agric. 2016;96(1):73–78. doi: 10.1002/jsfa.7061.

Matilla MA, Krell T. The effect of bacterial chemotaxis on host infection and pathogenicity. FEMS Microbiol Rev. 2018;42(1). doi: 10.1093/femsre/fux052.

Matsuoka Y, Li X, Bennett V. Adducin: structure, function and regulation. Cell Mol Life Sci. 2000;57(6):884–895. doi: 10.1007/PL00000731.

Mauck KE, Chesnais Q, Shapiro LR. Evolutionary Determinants of Host and Vector Manipulation by Plant Viruses. Adv Virus Res. 2018;101:189–250. doi: 10.1016/bs.aivir.2018.02.007.

McFarland L, Francetić O, Kumamoto CA. A mutation of *Escherichia coli* SecA protein that partially compensates for the absence of SecB. J Bacteriol. 1993;175(8):2255–2262. doi: 10.1128/jb.175.8.2255-2262.1993.

Melcarne C, Lemaitre B, Kurant E. Phagocytosis in Drosophila: From molecules and cellular machinery to physiology. Insect Biochem Mol Biol. 2019;109:1–12. doi: 10.1016/j.ibmb.2019.04.002.

Merz F, Hoffmann A, Rutkowska A, Zachmann-Brand B, Bukau B, Deuerling E. The C-terminal domain of *Escherichia coli* trigger factor represents the central module of its chaperone activity. J Biol Chem. 2006;281(42):31963–31971. doi: 10.1074/jbc.M605164200.

Moens S, Vanderleyden J. Functions of bacterial flagella. Crit Rev Microbiol. 1996;22(2):67–100. doi: 10.3109/10408419609106456.

Molki B, Thi Ha P, Mohamed A, Killiny N, Gang DR, Omsland A, Beyenal H. Physiochemical changes mediated by “*Candidatus* Liberibacter asiaticus” in Asian citrus psyllids. Sci Rep. 2019;9(1):16375. doi: 10.1038/s41598-019-52692-7.

Moy RH, Cherry S. Antimicrobial autophagy: a conserved innate immune response in Drosophila. J Innate Immun. 2013;5(5):444–455. doi: 10.1159/000350326.

Münch D, Amdam GV, Wolschin F. Ageing in a eusocial insect: molecular and physiological characteristics of life span plasticity in the honey bee. Funct Ecol. 2008;22(3):407–421. doi: 10.1111/j.1365-2435.2008.01419.x.

Nakhleh J, Moussawi LE, Osta MA. The melanization response in insect immunity. Adv In Insect Phys. 2017;52:83–109. doi: 10.1016/bs.aiip.2016.11.002.

Nardi JB, Miller LA, Bee CM. Interfaces between microbes and membranes of host epithelial cells in hemipteran midguts. J Morphol. 2019;280(7):1046–1060. doi: 10.1002/jmor.21000.

Neish AS. Redox signaling mediated by the gut microbiota. Free Radic Res. 2013;47(11):950–7. doi: 10.3109/10715762.2013.833331.

Norville IH, Breitbach K, Eske-Pogodda K, Harmer NJ, Sarkar-Tyson M, Titball RW, Steinmetz I. A novel FK-506-binding-like protein that lacks peptidyl-prolyl isomerase activity is involved in intracellular infection and in vivo virulence of *Burkholderia pseudomallei*. Microbiology. 2011;157:2629–2638. doi: 10.1099/mic.0.049163-0.

Pandey S, Tripathi D, Khubaib M, Kumar A, Sheikh JA, Sumanlatha G, Ehtesham NZ, Hasnain SE. *Mycobacterium tuberculosis* Peptidyl-Prolyl Isomerases Are Immunogenic, Alter Cytokine Profile and Aid in Intracellular Survival. Front Cell Infect Microbiol. 2017;7:38. doi: 10.3389/fcimb.2017.00038.

Panz M, Vitos-Faleato J, Jendretzki A, Heinisch JJ, Paululat A, Meyer H. A novel role for the non-catalytic intracellular domain of Neprilysins in muscle physiology. Biol Cell. 2012;104(9):553–568. doi: 10.1111/boc.201100069.

Paone P, Cani PD. Mucus barrier, mucins and gut microbiota: the expected slimy partners? Gut. 2020;69(12):2232–2243. doi: 10.1136/gutjnl-2020-322260.

Perilla-Henao LM, Casteel CL. Vector-Borne Bacterial Plant Pathogens: Interactions with Hemipteran Insects and Plants. Front Plant Sci. 2016 Aug 9;7:1163. doi: 10.3389/fpls.2016.01163.

Perkins A, Nelson KJ, Parsonage D, Poole LB, Karplus PA. Peroxiredoxins: guardians against oxidative stress and modulators of peroxide signaling. Trends Biochem Sci. 2015;40(8):435–445. doi: 10.1016/j.tibs.2015.05.001.

Porter ME. Axonemal dyneins: assembly, organization, and regulation. Curr Opin Cell Biol. 1996;8(1):10–7. doi: 10.1016/s0955-0674(96)80042-1.

Prasad S, Xu J, Zhang Y, Wang N. SEC-Translocon Dependent Extracytoplasmic Proteins of *Candidatus* Liberibacter asiaticus. Front Microbiol. 2016;7:1989. doi: 10.3389/fmicb.2016.01989.

Quigley EM. Gut bacteria in health and disease. Gastroenterol Hepatol (N Y). 2013;9(9):560–9.

Quintana-Hayashi MP, Mahu M, De Pauw N, Boyen F, Pasmans F, Martel A, et al. The levels of *Brachyspira hyodysenteriae* binding to porcine colonic mucins differ between individuals, and binding is increased to mucins from infected pigs with *de novo* MUC5AC synthesis. Infect Immun. 2015;83(4):1610–9. doi: 10.1128/IAI.03073-14.

Rahman MM, Franch-Marro X, Maestro JL, Martin D, Casali A. Local Juvenile Hormone activity regulates gut homeostasis and tumor growth in adult *Drosophila*. Sci Rep. 2017;7(1):11677. doi: 10.1038/s41598-017-11199-9.

Ramsey JS, Chavez JD, Johnson R, Hosseinzadeh S, Mahoney JE, Mohr JP, et al. Protein interaction networks at the host-microbe interface in *Diaphorina citri*, the insect vector of the citrus greening pathogen. R Soc Open Sci. 2017;4(2):160545. doi: 10.1098/rsos.160545.

Ramsey JS, Johnson RS, Hoki JS, Kruse A, Mahoney J, Hilf ME, et al. Metabolic Interplay between the Asian Citrus Psyllid and Its *Profftella* Symbiont: An Achilles’ Heel of the Citrus Greening Insect Vector. PLoS One. 2015;10(11):e0140826. doi: 10.1371/journal.pone.0140826.

Rasch J, Ünal CM, Klages A, Karsli Ü, Heinsohn N, Brouwer RMHJ, et al. Peptidyl-Prolyl-cis/trans-Isomerases Mip and PpiB of *Legionella pneumophila* Contribute to Surface Translocation, Growth at Suboptimal Temperature, and Infection. Infect Immun. 2018;87(1):e00939–17. doi: 10.1128/IAI.00939-17.

Ray S, Da Costa R, Thakur S, Nandi D. *Salmonella* Typhimurium encoded cold shock protein E is essential for motility and biofilm formation. Microbiology. 2020;166(5):460–473. doi: 10.1099/mic.0.000900.

Redder P, Hausmann S, Khemici V, Yasrebi H, Linder P. Bacterial versatility requires DEAD-box RNA helicases. FEMS Microbiol Rev. 2015;39(3):392–412. doi: 10.1093/femsre/fuv011.

Riddell CE, Lobaton Garces JD, Adams S, Barribeau SM, Twell D, Mallon EB. Differential gene expression and alternative splicing in insect immune specificity. BMC Genomics. 2014;15(1):1031. doi: 10.1186/1471-2164-15-1031.

Roberts IS. The biochemistry and genetics of capsular polysaccharide production in bacteria. Annu Rev Microbiol. 1996;50:285–315. doi: 10.1146/annurev.micro.50.1.285.

Robinson CG, Roy CR. Attachment and fusion of endoplasmic reticulum with vacuoles containing *Legionella pneumophila*. Cell Microbiol. 2006;8(5):793–805. doi: 10.1111/j.1462-5822.2005.00666.x.

Roche B, Aussel L, Ezraty B, Mandin P, Py B, Barras F. Iron/sulfur proteins biogenesis in prokaryotes: formation, regulation and diversity. Biochim Biophys Acta. 2013;1827(3):455–469. doi: 10.1016/j.bbabio.2012.12.010.

Rothman JE. Mechanisms of intracellular protein transport. Nature. 1994;372(6501):55–63. doi: 10.1038/372055a0.

Santos CA, Janissen R, Toledo MA, Beloti LL, Azzoni AR, Cotta MA, Souza AP. Characterization of the TolB-Pal trans-envelope complex from *Xylella fastidiosa* reveals a dynamic and coordinated protein expression profile during the biofilm development process. Biochim Biophys Acta. 2015;1854:1372–1381. doi: 10.1016/j.bbapap.2015.05.018.

Santos-Beneit F. The Pho regulon: a huge regulatory network in bacteria. Front Microbiol. 2015;6:402. doi: 10.3389/fmicb.2015.00402.

Santos-Ortega Y, Killiny N. Silencing of sucrose hydrolase causes nymph mortality and disturbs adult osmotic homeostasis in *Diaphorina citri* (Hemiptera: Liviidae). Insect Biochem Mol Biol. 2018;101:131–143. doi: 10.1016/j.ibmb.2018.09.003.

Schroeder BO. Fight them or feed them: how the intestinal mucus layer manages the gut microbiota. Gastroenterol Rep (Oxf). 2019;7(1):3–12. doi: 10.1093/gastro/goy052.

Shears SB, Hayakawa Y. Functional Multiplicity of an Insect Cytokine Family Assists Defense Against Environmental Stress. Front Physiol. 2019;10:222. doi: 10.3389/fphys.2019.00222.

Silva CP, Silva JR, Vasconcelos FF, Petretski MD, Damatta RA, Ribeiro AF, Terra WR. Occurrence of midgut perimicrovillar membranes in paraneopteran insect orders with comments on their function and evolutionary significance. Arthropod Struct Dev. 2004;33(2):139–148. doi: 10.1016/j.asd.2003.12.002.

Singerman A, Rogers ME. The economic challenges of dealing with citrus greening: the case of Florida. J Integr Pest Manag. 2020;11(1):1–7. doi: 10.1093/jipm/pmz037.

Solari P, Rivelli N, De Rose F, Picciau L, Murru L, Stoffolano JG Jr, Liscia A. Opposite effects of 5-HT/AKH and octopamine on the crop contractions in adult *Drosophila melanogaster*: Evidence of a double brain-gut serotonergic circuitry. PLoS One. 2017;12(3):e0174172. doi: 10.1371/journal.pone.0174172.

Sohlenkamp C, López-Lara IM, Geiger O. Biosynthesis of phosphatidylcholine in bacteria. Prog Lipid Res. 2003;42(2):115–62. doi: 10.1016/s0163-7827(02)00050-4.

Stenbeck G. Soluble NSF-attachment proteins. Int J Biochem Cell Biol. 1998;30(5):573–7. doi: 10.1016/s1357-2725(97)00064-2.

Stokes BA, Yadav S, Shokal U, Smith LC, Eleftherianos I. Bacterial and fungal pattern recognition receptors in homologous innate signaling pathways of insects and mammals. Front Microbiol. 2015;6:19. doi: 10.3389/fmicb.2015.00019.

Strand M, Micchelli CA. Quiescent gastric stem cells maintain the adult *Drosophila* stomach. Proc Natl Acad Sci U S A. 2011;108(43):17696–701. doi: 10.1073/pnas.1109794108.

Thapa SP, De Francesco A, Trinh J, Gurung FB, Pang Z, Vidalakis G, Wang N, Ancona V, Ma W, Coaker G. Genome-wide analyses of Liberibacter species provides insights into evolution, phylogenetic relationships, and virulence factors. Mol Plant Pathol. 2020;21(5):716–731. doi: 10.1111/mpp.12925.

Theodorou MC, Theodorou EC, Kyriakidis DA. Involvement of AtoSC two-component system in *Escherichia coli* flagellar regulon. Amino Acids. 2012;43(2):833–844. doi: 10.1007/s00726-011-1140-7.

Tjota M, Lee SK, Wu J, Williams JA, Khanna MR, Thomas GH. Annexin B9 binds to β(H)-spectrin and is required for multivesicular body function in *Drosophila*. J Cell Sci. 2011;124:2914–2926. doi: 10.1242/jcs.078667.

Tomaseto AF, Marques RN, Fereres A, Zanardi OZ, Volpe HXL, Alquézar B, Peña L, Miranda MP. Orange jasmine as a trap crop to control *Diaphorina citri*. Sci Rep. 2019;9(1):2070. doi: 10.1038/s41598-019-38597-5.

Treutter D. Significance of flavonoids in plant resistance: a review. Environ Chem Lett. 2006;4:147–157. doi: 10.1007/s10311-006-0068-8.

Turner AJ, Isaac RE, Coates D. The neprilysin (NEP) family of zinc metalloendopeptidases: genomics and function. Bioessays. 2001;23(3):261–269. doi: 10.1002/1521-1878(200103)23:3<261::AID-BIES1036>3.0.CO;2-K.

Vallet-Gely I, Lemaitre B, Boccard F. Bacterial strategies to overcome insect defences. Nat Rev Microbiol. 2008;6(4):302–313. doi: 10.1038/nrmicro1870.

Vande Velde C, Cizeau J, Dubik D, Alimonti J, Brown T, Israels S, et al. BNIP3 and genetic control of necrosis-like cell death through the mitochondrial permeability transition pore. Mol Cell Biol. 2000;20(15):5454–5468. doi: 10.1128/mcb.20.15.5454-5468.2000.

Wang C, Chen Y, Zhou H, Li X, Tan Z. Adaptation mechanisms of *Rhodococcus* sp. CNS16 under different temperature gradients: Physiological and transcriptome. Chemosphere. 2020;238:124571. doi: 10.1016/j.chemosphere.2019.124571.

Wang N, Pierson EA, Setubal JC, Xu J, Levy JG, Zhang Y, Li J, Rangel LT, Martins J Jr. The *Candidatus* Liberibacter-Host Interface: Insights into Pathogenesis Mechanisms and Disease Control. Annu Rev Phytopathol. 2017;55:451–482. doi: 10.1146/annurev-phyto-080516-035513.

Webb BA, Strand MR, Dickey SE, Beck MH, Hilgarth RS, Barney WE, et al. Polydnavirus genomes reflect their dual roles as mutualists and pathogens. Virology. 2006;347(1):160–74. doi: 10.1016/j.virol.2005.11.010.

Wegener C, Veenstra JA. Chemical identity, function and regulation of enteroendocrine peptides in insects. Curr Opin Insect Sci. 2015;11:8–13. doi: 10.1016/j.cois.2015.07.003.

Weinbauer MG. Ecology of prokaryotic viruses. FEMS Microbiol Rev. 2004;28(2):127–81. doi: 10.1016/j.femsre.2003.08.001.

Witke W. The role of profilin complexes in cell motility and other cellular processes. Trends Cell Biol. 2004;14(8):461–469. doi: 10.1016/j.tcb.2004.07.003.

Wu K, Li S, Wang J, Ni Y, Huang W, Liu Q, Ling E. Peptide Hormones in the Insect Midgut. Front Physiol. 2020;11:191. doi: 10.3389/fphys.2020.00191.

Wu P, Sun P, Nie K, Zhu Y, Shi M, Xiao C, et al. A Gut Commensal Bacterium Promotes Mosquito Permissiveness to Arboviruses. Cell Host Microbe. 2019;25(1):101–112.e5. doi: 10.1016/j.chom.2018.11.004.

Wu SC, Liao CW, Pan RL, Juang JL. Infection-induced intestinal oxidative stress triggers organ-to-organ immunological communication in *Drosophila*. Cell Host Microbe. 2012;11(4):410–417. doi: 10.1016/j.chom.2012.03.004.

Wulff NA, Daniel B, Sassi RS, Moreira AS, Bassanezi RB, Sala I, et al. Incidence of *Diaphorina citri* Carrying *Candidatus* Liberibacter asiaticus in Brazil’s Citrus Belt. Insects. 2020;11(10):672. doi: 10.3390/insects11100672.

Yu HZ, Li NY, Zeng XD, Song JC, Yu XD, Su HN, et al. Transcriptome Analyses of *Diaphorina citri* Midgut Responses to *Candidatus* Liberibacter Asiaticus Infection. Insects. 2020;11(3):171. doi: 10.3390/insects11030171.

Yu X, Killiny N. RNA interference-mediated control of Asian citrus psyllid, the vector of the huanglongbing bacterial pathogen. Trop Plant Pathol. 2020;45:298–305. doi: 10.1007/s40858-020-00356-7.

Zalewska M, Kochman A, Estève JP, Lopez F, Chaoui K, Susini C, Ozyhar A, Kochman M. Juvenile hormone binding protein traffic - Interaction with ATP synthase and lipid transfer proteins. Biochim Biophys Acta. 2009;1788(9):1695–705. doi: 10.1016/j.bbamem.2009.04.022.

Zhang B, Gong J, Zhang W, Xiao R, Liu J, Xu XZS. Brain-gut communications via distinct neuroendocrine signals bidirectionally regulate longevity in *C. elegans*. Genes Dev. 2018;32(3-4):258–270. doi: 10.1101/gad.309625.117.

Zhao M, Wang S, Li F, Dong D, Wu B. Arylsulfatase B Mediates the Sulfonation-Transport Interplay in Human Embryonic Kidney 293 Cells Overexpressing Sulfotransferase 1A3. Drug Metab Dispos. 2016;44(9):1441–1449. doi: 10.1124/dmd.116.070938

